# Building Brains for Robots: A Hands-On Approach to Learning Neuroscience in the Classroom

**DOI:** 10.1101/2024.05.15.594177

**Authors:** Raha Kannan, Maribel Gendreau, Alex Hatch, Sydney K. Free, Kithinji Muriungi, Yash A. Garje, Jennifer DeBoer, Gregory J. Gage, Christopher A. Harris

## Abstract

As the relevance of neuroscience in education grows, effective methods for teaching this complex subject in high school classrooms remain elusive. Integrating classroom experiments with brain-based robots offers a promising solution. This paper presents a structured curriculum designed around the use of camera-equipped mobile robots which enables students to construct and explore artificial neural networks. Through this hands-on approach, students engage directly with core concepts in neuroscience, learning to model spiking neural networks, decision-making processes in the basal ganglia, and principles of learning and memory. The curriculum not only makes challenging neuroscience concepts accessible and engaging but also demonstrates significant improvements in students’ understanding and self-efficacy. By detailing the curriculum’s development, implementation, and educational outcomes, this study outlines a scalable model for incorporating advanced scientific topics into secondary education, paving the way for a deeper student understanding of both theoretical neuroscience and its practical applications.

## Introduction

In the modern educational landscape, understanding the brain’s complexities is not just beneficial - it’s essential. The prevalence of brain disorders across the globe underscores the urgent need for a new generation of scientists, physicians and engineers who are adept in neuroscience. Moreover, the proliferation of neural network-based technologies in our daily lives - from artificial intelligence (AI) to autonomous systems - demands that formal educational curricula evolve to include these advanced concepts. Neuroscience is not yet part of most education standards, but there is a growing movement to include it (Gage, 2019; Minen et al., 2023). While student laboratories with biological nervous system activities and preparations are gradually finding their way into secondary education, there is a dearth of technologies and curricula for experimenting with computational models of neural networks. Integrating neural network modeling into education prepares students for an AI-powered future, deepens their understanding of the brain’s biology, and enhances their potential to advance technology and treat neurological disorders.

To meet these educational demands, we have developed a program for high schoolers that combines computational neuroscience and neural network modeling with hands-on activities involving camera-equipped mobile robots, called SpikerBots (Figure 1). Through a dedicated app, students craft neural networks for the robots, exploring various neurons, synapses, and behaviors. This method aligns with and applies Albert Bandura’s Social Cognitive Theory, which views learning as a social endeavor influenced by the dynamic between individuals, their environment, and their own actions, highlighting the importance of self-efficacy—or confidence in one’s abilities for specific tasks (Bandura, 1986). Our initiative offers broadly applicable potential for teachers and learners, especially in secondary school settings, as it seeks to bolster students’ self-efficacy by engaging them in interactive experiments with SpikerBots, fostering learning through exploration and active participation. In teams, students hypothesize, build, and refine neural networks to achieve desired behaviors. The curriculum offers flexibility in achieving outcomes (multiple networks can achieve the same behavior) while providing structured guidance for teachers and support for students. Piloted in several high schools, the program has received positive evaluations from both students and educators, and has demonstrated significantly enhanced neuroscience conceptual understanding and self-efficacy post-intervention (Harris et al., 2020).

**Figure 1.**
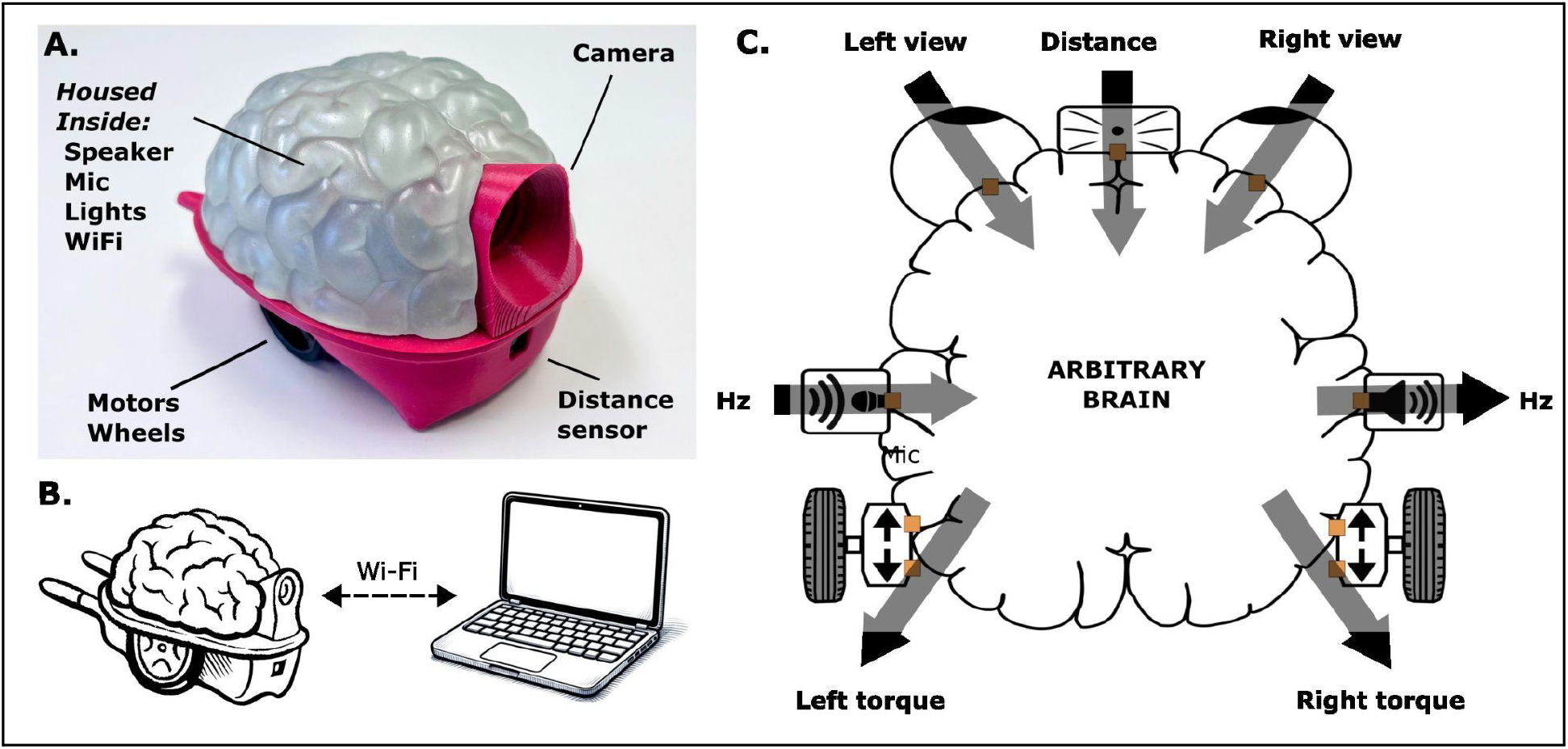
A robot for neuroscience education. The SpikerBot robot **(A)** is a mobile robot designed for the high school science classroom. It maintains a permanent WiFi connection with the SpikerBot app **(B)**, which runs its neural networks. A brain-shaped interface **(C)** is used to visualize and edit the robot’s neural networks.

A survey of 84 secondary school science teachers revealed an interest in incorporating SpikerBots into their teaching (deFreitas et al., 2021). However, the same study also pointed to a need for a well-structured curriculum that would enable teachers to independently use robot-based neuroscience instruction in their classrooms, and address fundamental questions: Which neural networks should students be challenged to construct? What instructions and tools do students need to successfully build brains with desired behavioral output? How should lessons be arranged to promote teamwork and the development of self-efficacy?

Here, we present a brief report with clear potential for pushing the boundaries of formal and informal inquiry-based learning. Our study pilots and tests the impact of a robot-based neuroscience curriculum for high schools, comprising two main components:

- **3 Lesson Plans**. Each lesson plan includes learning objectives, brain design exercises, step-by-step solutions, and student handouts (Supplemental Materials 1-3).
- **Primer for Teachers**. This guide provides teachers with foundational neuroscience knowledge, SpikerBot use instructions, and best practices for teaching with SpikerBots in the classroom (Supplemental Materials 4).

The lessons introduce three complementary approaches to understanding and designing brains:

- **Lesson 1**. Students learn how spiking neural networks generate behavior by manually connecting neurons and synapses to the robot’s sensors and effectors.
- **Lesson 2**. Students learn how the brain makes choices by constructing a model of the basal ganglia, a brain region involved in decision-making.
- **Lesson 3**. Students collect data and train neural networks to recognize patterns and achieve goals.

In the next section, we describe the curriculum in detail. Then we report the results of a 1-week pilot enactment of the curriculum at a large suburban public high school (n = 26 students). We end by discussing lessons learned and future directions.

### Curriculum Structure and Learning Objectives

Our approach combines a mobile robot (Mondada et al., 2017; Tani et al., 2017; Paull et al., 2017) with a pointer-based software application that lets students create and simulate spiking neural networks (Dragly et al., 2017; Tsai & Yoshimi, 2023). Students develop a foundational understanding of the link between structure and function in the nervous system through direct observation and manipulation of the robot’s neural networks.

The curriculum focuses on three core concepts in computational neuroscience: spiking neural networks, the basal ganglia, and learning. Spiking neurons and synapses are the brain’s basic building blocks, enabling a vast range of behaviors. The basal ganglia, a deep brain structure, is crucial in choosing among these available behaviors. Lastly, the remarkable ability of neural networks to learn empowers them to understand and interact with the world around them.

The curriculum is designed to enhance hands-on learning by emphasizing active student participation. Lessons start with the teacher presenting important neuroscience ideas and showing how different neural network models result in specific robot actions. After this introduction, students independently tackle the exercises provided in the lesson, promoting an engaged learning experience. The curriculum aims for a 25/75 balance between teacher instruction and active learning activities. Exercises are designed for small group work (2-5 students), with each group equipped with a SpikerBot robot, a laptop running the SpikerBot app, and lesson handouts.

### Lesson 1: Spiking Neural Networks Generate Behavior

#### Rationale

Lesson 1 is an introduction to fundamental computational neuroscience and neurorobotics. Students learn to construct their own networks of spiking neurons and synapses, interface them with their robot’s sensors and effectors, and evaluate the resulting behaviors. The lesson enables students to develop a strong conceptual understanding of the relationship between the structure of a neural network and its function (how it behaves).

#### Preparation

After sharing background information on biological and artificial neural networks, the teacher demonstrates to the students how to use the SpikerBot app to build and run neural networks. By the end of the demonstration, students should understand how to switch between the app’s three modes of operation (Main Menu, Runtime, and Design) and how to add, connect, edit, and delete individual neurons.

#### Exercises

The first exercise (L1E1, Figure 2A) tasks students with creating a brain that can avoid obstacles. One solution, provided to the teacher with assembly instructions in the lesson materials, involves a single neuron that detects nearby objects and drives the robot backward. By using just one neuron to perform a useful behavior, students gain an initial understanding of how sensory information can lead directly to action.

**Figure 2.**
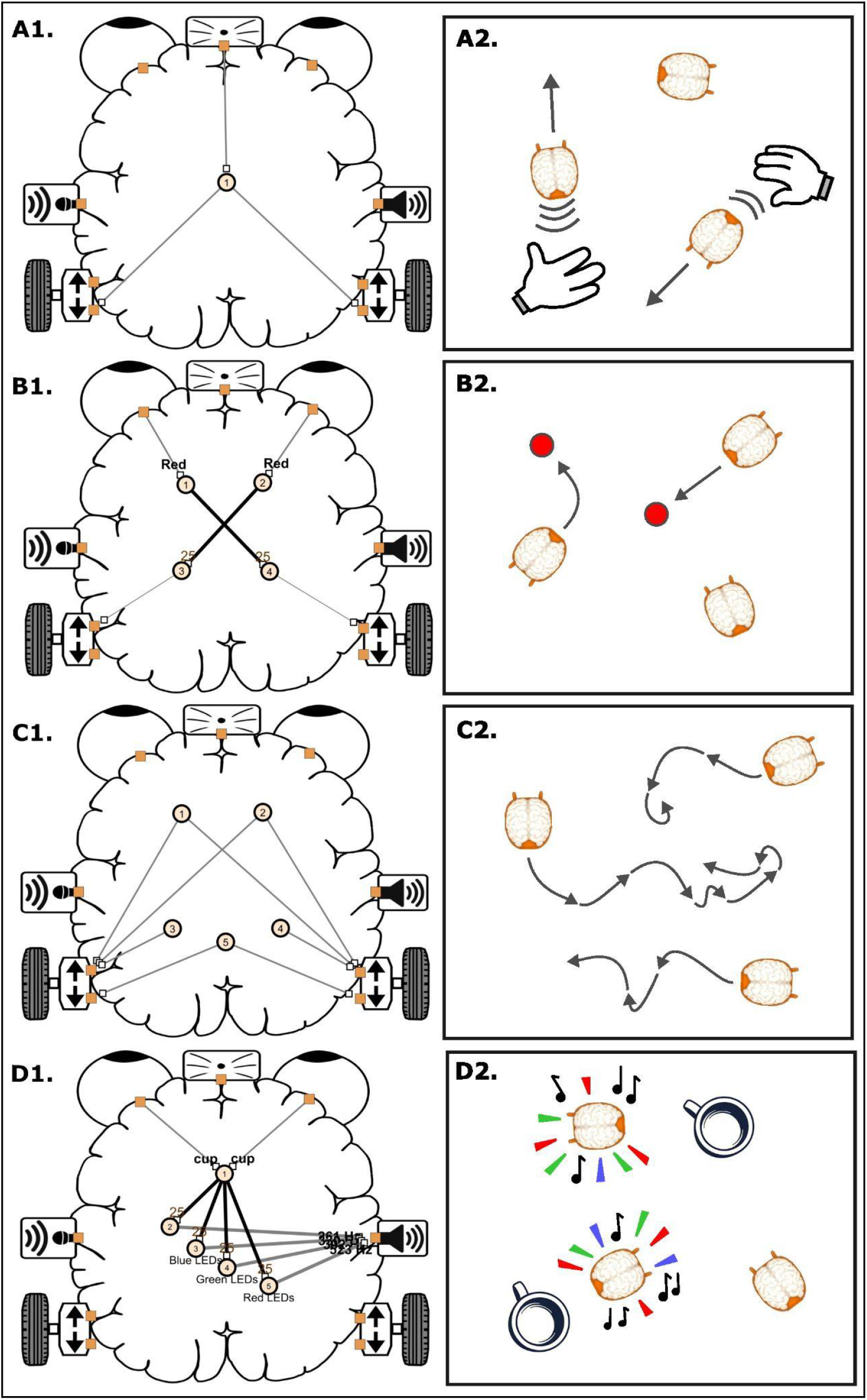
Lesson 1: Spiking Neural Networks. Students build brains expressing four different behaviors: obstacle avoidance **(A)**, color approach **(B)**, exploration **(C)**, and blinking and beeping at cups **(D)**.

The second exercise (L1E2, Figure 2B) is to design a brain that can approach a moving target. The provided solution employs four neurons: two that detect the color red on the left or right side of the visual field and two that drive the motor on the opposite side of the brain forward. By making the detection of red on the left side trigger forward motion on the right side, and vice versa, the brain effectively turns the robot to face the red target when it is off-center and makes the robot approach the target when it is directly ahead. The robot should even be able to follow moving red targets. The exercise shows how a small number of neurons can produce life-like, goal-directed behaviors (Braitenberg, 1984).

The third exercise (L1E3, Figure 2C) tasks students with designing a brain that explores its surroundings autonomously. This involves using neurons that are spontaneously active. The SpikerBot app implements the Izhikevich neuron model, which has two parameters that govern spike rate and burst duration, respectively (Izhikevich, 2003). These parameters are easily adjusted in the SpikerBot app to create neurons with unpredictable dynamics. By adding neurons that emit spontaneous bursts of spikes and connecting those neurons to the robot’s motors, students make the robot perform ‘random walks’, a life-like form of exploratory behavior.

In the lesson’s final exercise (L1E4, Figure 2D), students design a brain that blinks and beeps when it encounters coffee cups. To facilitate this, the SpikerBot app utilizes a large artificial neural network (GoogLeNet) that can recognize a wide range of objects (animals, fruits, tools, etc.) in image data. Students choose which objects individual neurons should respond to from a dropdown menu. The exercise’s provided solution employs a single cup-detecting neuron, which connects to four neurons that produce sounds and activate the robot’s LEDs. As students present the robot with different visual inputs, the app shows what the robot is seeing and how its brain is responding, creating an interactive demonstration of object recognition in neural networks.

By the end of Lesson 1, students will have a grasp of how simple neural networks can generate interesting behaviors, setting the stage for exploring more complex neuroscience concepts in subsequent lessons.

### Lesson 2: The Basal Ganglia Makes Decisions

#### Rationale

Decision-making in the brain involves selecting between diverse neural networks with different functions or behavioral outputs. This selection process is largely managed by a brain area known as the basal ganglia, which integrates (considers) inputs from multiple brain areas and selectively activates individual behavior-generating networks throughout the brain (Grillner and Robertson, 2016; Prescott et al., 2024). This is called a ‘winner-take-all’ selection process. In Lesson 2, students learn to build a computational model of the basal ganglia that can select the appropriate action at the correct time in order to complete a search task (finding hidden coffee cups) (Figure 3). The exercise aims to give students a practical understanding of how the brain makes smart decisions.

**Figure 3.**
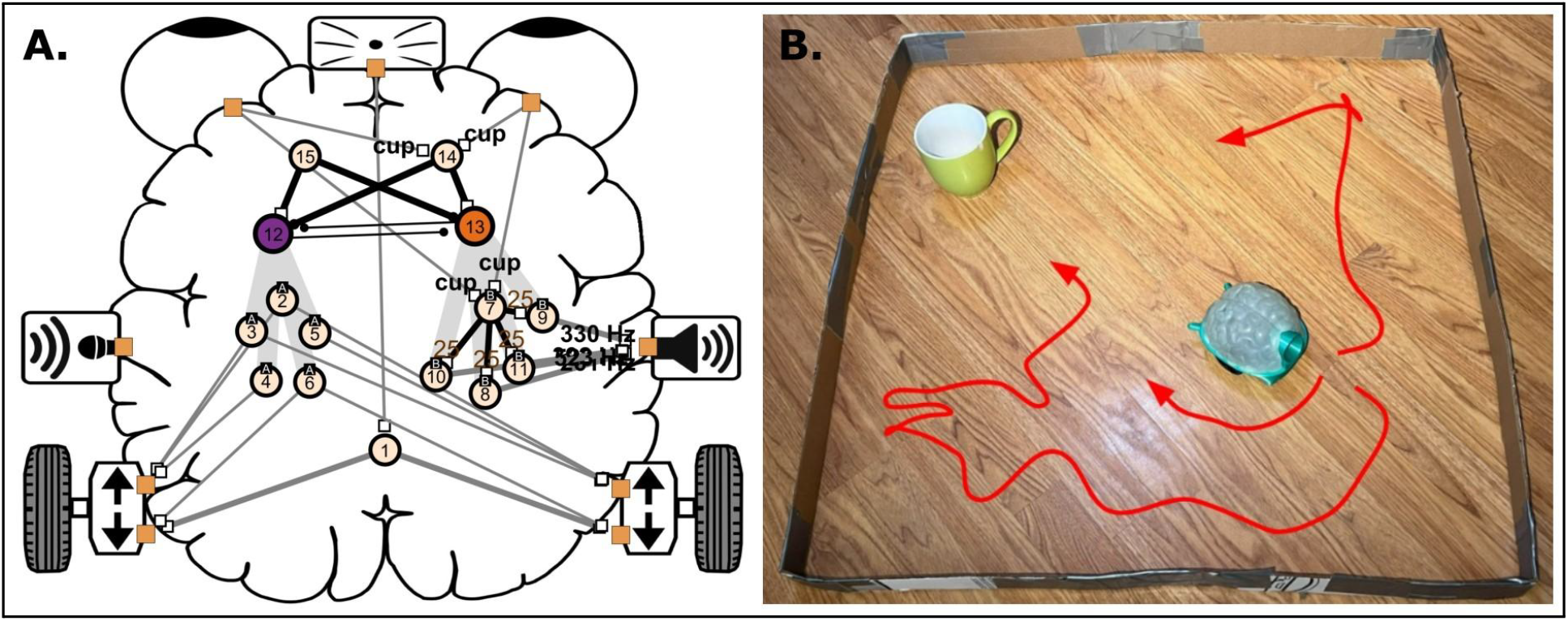
Lesson 2: Basal Ganglia. Students develop a brain **(A)** that can find a target (a cup) that is initially out of sight **(B)**.

#### Preparation

We recommend conducting the search task in an enclosed arena to allow for repeatable, timed experiments. The arena also allows students to work with the robots on tabletops rather than on the floor. To make an arena, cut cardboard strips (∼1 m by ∼5 cm) and tape them together to form a square (Figure 3B). In addition, each group needs access to a timer.

#### Exercises

In Lesson 2, students are tasked with creating a brain that can find one or more coffee cups that are initially out of sight (Figure 3B). To do this, students build a brain featuring a basal ganglia model that can choose to explore in the absence of cups and to blink and beep in the presence of cups. The construction of this complex brain is spread over two exercises.

In the first exercise (L2E1), students learn how to import previously saved networks. First, an obstacle-avoiding neuron (neuron 1 in Figure 3A) is added to the empty brain to stop it from colliding with the arena wall. Then the exploratory network L1E3 (neurons 2-6) and the cup-indicating network L1E4 (neurons 7-11) are added to the brain. The two networks are activated, one at a time, by the basal ganglia model, but without synaptic input to guide it, the basal ganglia will randomly select each network approximately 50% of the time regardless of whether a cup is in view. This makes the search behavior slow and ineffective.

In the second exercise (L2E2), students add ‘striatal’ neurons to improve the basal ganglia’s decision-making ability. Striatal neurons integrate signals from different parts of the brain to determine which action the basal ganglia will select at any given time (Cox and Witten, 2019). First, students assign a striatal neuron to each behavior (neurons 12-13). Then, students add a cup-detecting neuron (neuron 14) that inhibits the exploration-associated striatal neuron (neuron 12) and stimulates the cup indication-associated striatal neuron (neuron 14). This has the effect of biasing the basal ganglia’s selection process toward the indication behavior (blink and beep) in the presence of cups. Finally, students add a spontaneously active neuron (neuron 15) that stimulates exploration and inhibits indication via their respective striatal neurons. These synaptic connections bias the basal ganglia in favor of exploration in the absence of cups. The exercise teaches students how the vertebrate brain needs to be organized to make intelligent decisions, and allows them to combine multiple neural networks in a single brain.

### Lesson 3: Neural Networks Learn to Recognize Patterns and Achieve Goals

#### Rationale

Artificial neural networks have massively extended the abilities of AI in recent years by mimicking the brain’s method of learning through modification of synaptic connections between neurons (Hassabis et al., 2017; Zador et al., 2023). ‘Associative learning’ is when a brain or AI learns to identify patterns in data without being told ahead of time what the patterns are (e.g. learning to recognize a new face, voice, or location). ‘Reinforcement learning’ (RL) is when the brain or AI learns to select actions that lead to desired outcomes and avoid actions that lead to undesired outcomes (e.g. how to ride a bike or play an instrument). The SpikerBot app leverages machine-learning algorithms to demystify associative and reinforcement learning for students (Figure 4). While there are differences between machine learning and the learning that goes on in brains, the outcomes are comparable: neural networks learn to recognize patterns and achieve goals. Lesson 3 teaches students how to train neural networks to find cups in the cardboard arena significantly faster than the brain developed in Lesson 2.

**Figure 4.**
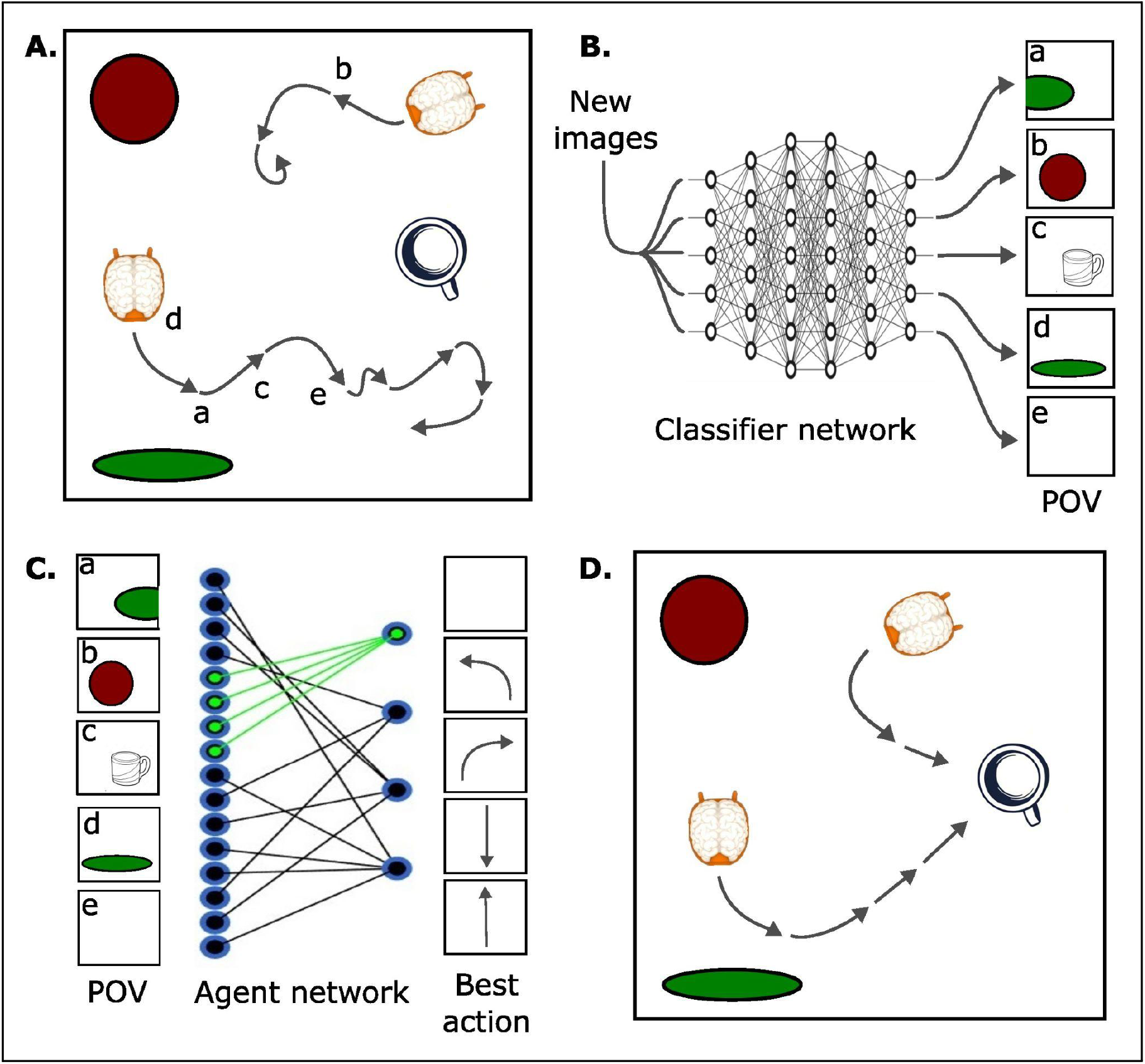
Lesson 3: Learning. Students use machine learning to train neural networks to navigate new environments. First, robots explore randomly **(A)**, saving camera frames and motor commands. Then, a classifier network **(B)** is trained to recognize recurring points of view (POVs). Finally, an agent network **(C)** is trained to navigate to a specific POV (chosen by the students, indicated by green lines) from anywhere in the new environment. The trained network’s ability to reach the goal POV in the real world is evaluated **(D)**.

#### Preparation

Machine learning requires a lot of training data. In Lesson 3, the training data consists of camera images and motor commands recorded by the SpikerBot as it explores the cardboard arena (Figure 4A). Combining the obstacle avoiding brain L1E1 with the exploring brain L1E3 creates a good explorer for this purpose. The data needs to be recorded and processed before the start of Lesson 3. We find that 30 min of exploration data is sufficient to train a neural network to find cups in the arena. However, more data will produce better results.

The learning process is entirely automated. It begins by identifying about 20 categories of images with similar visual features (Figure 4B). Each category corresponds to the camera view from a specific location in the arena (Figure 4C). It then trains a classifier network (a large neural network consisting of thousands of neurons) to categorize new images. Next, the training process generates a model (transition matrix) that estimates how likely different actions (motor outputs) are to move the robot from any one location to any other. Finally, a 2-layer neural network called an ‘agent’ (Figure 4D) is trained to navigate from any given location in the arena to one where the cups are in view. The entire learning process takes about 7 hrs per 1 hour of exploration data on a laptop with an i5 CPU and integrated graphics.

#### Exercises

In the first exercise (L3E1), students are invited to load the trained location classifier network and test it by physically placing the robot in different locations in the arena and confirming that specific neurons are consistently activated in the brain. This exercise gives students an intuition for how well the neural network has learned the new environment.

In the second exercise (L3E2), students identify cup-containing locations / points of view and train an ‘agent’ network to navigate to them. This involves a few mouse clicks and ∼10 min of training. Alternatively, the agent network can be trained by the teacher before the lesson starts. Students then load the trained classifier network with the trained agent network added as a final layer, such that every location in the arena triggers a specific action (combination of motor torques) that eventually results in the robot finding the target (Figure 4D). The exercise requires students to evaluate their trained brain’s ability to find the cups in the arena by timing it and comparing its performance to that of a randomly exploring brain (L2E2). In our tests, the trained brain is more than twice as fast.

### Pilot Workshop

We conducted a pilot workshop at Novi High School, a large, ethnically diverse public suburban high school in Michigan, to assess our curriculum via a structured five-day program, dedicating one hour daily. Our protocol was approved by the Purdue University IRB (September 25, 2023, protocol number IRB-2023-1376). Twenty-six students (ages 16-18) participated in the workshop as part of their regular AP (Advance Placement) biology instruction. Students worked in teams of 2-5 per robot-laptop setup. Each team was provided with a laptop and a SpikerBot.

On day 1, students took the pre-workshop quiz and started learning to use the app. On day 2, students engaged directly with the robots to complete the Lesson 1 exercises. Due to time constraints, some students used pre-built brains instead of creating their own. On day 3, students worked with basal ganglia models to generate effective cup-finding behavior. Training data for deep learning was collected during the lesson and processed overnight. On days 4 and 5, students learned to import trained networks and evaluate their ability to find cups. Some groups explored changing the firing rates of different neurons to improve overall performance. We ended with a post-workshop quiz and debrief.

The workshop yielded notable advancements in the students’ understanding of the different topics, paralleling a significant increase in self-efficacy (belief in one’s ability to complete a task).

The pre-and post-workshop quizzes used one open-ended question for each of the three lessons to evaluate neuroscience content learning outcomes:

- Q1: Describe three types of neurons with different spiking patterns
- Q2: How does the basal ganglia control behavior?
- Q3: Describe how neural networks learn new things

Answers were scored 1-3 (Incorrect, Partial, Correct) and tabulated (Table 1). The number of incorrect answers decreased in the post-workshop survey, whereas the number of partial and correct answers increased. Fisher’s exact test showed significant changes on all three questions.

**Table 1.** Conceptual learning results.

|  | Pre |  |  | Post |  |  | p-value |
| --- | --- | --- | --- | --- | --- | --- | --- |
|  | Incorrect | Partial | Correct | Incorrect | Partial | Correct |  |
| **Q1** | 23 | 3 | 0 | 8 | 9 | 7 | $p < 0.01$ |
| **Q2** | 22 | 3 | 1 | 9 | 3 | 12 | $p < 0.01$ |
| **Q3** | 17 | 8 | 1 | 7 | 12 | 5 | 0.03 |

**Table 2.** Student attitude results.

|  | Pre | Post | Difference | p-value | Effect size |
| --- | --- | --- | --- | --- | --- |
| **Self-Efficacy** | 4.01 ± 1.43 | 2.77 ± 0.95 | Decrease of 1.24 | p < 0.01 | 0.846 |
| **Self-Concept** | 2.69 ± 0.93 | 2.61 ± 0.95 | Decrease of 0.08 | p > 0.05 | 0.180 |

The pre- and post-workshop surveys used Likert-scale questions to evaluate self-efficacy and self-concept (e.g., “I am confident I can understand complex material in neuroscience”). The complete list of questions is available in the Supplemental Materials. Students’ answers were reverse coded: a lower numerical score denotes higher self-efficacy values. We observed a decrease from 4.01 to 2.77 in self-efficacy scores in the pre-vs post-workshop survey, indicating significant improvement. Fisher’s exact test confirmed that the change was statistically significant. A small improvement in self-concept was not statistically significant.

These results show that our curriculum supports learning in three different areas of computational neuroscience at the high school level. However, more time and support must be dedicated to each lesson to ensure students can attempt all exercises and conclude the workshop with significant increases in confidence (self-efficacy) and beliefs about themselves (self-concept).

## Discussion

We have developed an innovative high school curriculum that seamlessly integrates robotics with foundational concepts in neuroscience and neural modeling. The pilot workshop at Novi High School demonstrated substantive and statistically significant improvements in students’ understanding of computational neuroscience concepts and in self-efficacy to perform in neuroscience. Engaging with SpikerBots allowed students to experience firsthand the dynamic interplay between their actions, the robots’ responses, and the learning environment, fostering a sense of achievement and bolstering their confidence. Such hands-on activities not only deepened their conceptual understanding but also significantly reinforced their belief in their ability to execute complex tasks effectively—a central pillar of the construct of self-efficacy. However, we also note that we did not definitively identify a change in students’ self-concept. These contrasting results fit with the conceptual distinction between self-efficacy (confidence to perform a given task) and self-concept (perception of oneself, in this case, in the field of science). Prior research concurs that, while there is a positive relationship between the two constructs (Arens et al., 2020), self-efficacy is more strongly and immediately affected by inquiry-based learning opportunities (Jansen et al., 2015). Self-concept, on the other hand, is much more relevant to peer group comparisons and may take a longer time to change (e.g. Ehmke et al., 2010, Jansen et al., 2015); in our case, since the whole class was struggling through issues with the neurorobots together, there was likely not enough time or discrimination in success to foster change in self-concept. However, future work building on our report’s example could further focus on self-concept by providing longer-term interventions, and self-concept is, crucially, more closely related to future aspirations.

The importance of active learning in enhancing students’ self-efficacy, self-concept, and conceptual understanding cannot be overstated. Building on the groundwork of Bandura’s motivational theory, numerous studies have robustly demonstrated the profound impact of inquiry-based learning and authentic hands-on experiments on boosting students’ self-efficacy, science identity, and retention across STEM fields and diverse demographic groups (e.g., Swan et al., 2018, Ahmad et al., 2021, Jeskova et al., 2022, Wilczek et al. 2022, Sperling et al., 2024, Broder et al., 2019, Indorf et al., 2019). These studies lend robust support to our empirical observations, highlighting the critical importance of integrating real-world experimentation and inquiry-driven methods to significantly enhance students’ conceptual comprehension and sense of confidence in their ability to perform experiments and apply these concepts.

The principal challenge we encountered was time allocation, which limited our ability to delve deeply into the curriculum’s complex content. This issue was compounded by hardware and software difficulties, though we anticipate resolving these in future iterations. Analysis of video recordings from the workshop revealed that logistical challenges, such as pairing robots with computers and establishing WiFi connections, significantly affected student engagement. Often, only one student per group was actively engaged, with the rest observing, highlighting the need for more inclusive and interactive teaching methods. Despite these challenges, we noted an increase in engagement and emotional responses when teachers interacted directly with each group, suggesting that personal engagement and story-sharing are powerful tools for enhancing interest and confidence in the subject matter.

Our experience underscores the necessity of direct teacher-student interactions and additional time for students to complete all design exercises. Facing limitations in their knowledge, students did not significantly alter their perception of themselves as scientists. Nonetheless, the practical navigation through lessons in small groups and as a class effectively bolstered their confidence in performing relevant tasks.

In future workshops, we plan to emphasize clearer instructions and checklists for technical setup and allocate more time for each lesson. We will also enhance our use of classroom video monitoring to assess student engagement more accurately.

## Conclusion

This study highlights the potential of a robot-based curriculum to enrich students’ understanding of neuroscience, spiking neural networks, functional neuroanatomy, and artificial intelligence through a blend of theory, hands-on experimentation, and personal engagement. Our foray into neuroscience education through robotics marks a critical step toward more immersive and effective teaching methods. By confronting these challenges and capitalizing on our successes, we aim to inspire curiosity, enhance understanding, and foster innovation among neuroscience students. As we look ahead, the opportunity to further embed neuroscience into the STEM education framework is vast, promising a future where students can fully appreciate the complexity and marvel of the human brain.

## Supporting information

Lesson 1

Lesson 2

Lesson 3

Primer for Teachers

Survey

## Declarations

### Availability of data and materials

The datasets generated for this study are available on request to the corresponding authors.

### Authors’ contributions

RK and CAH conceived of the idea. RK, MG, AH, SKF and CAH developed the curriculum. CAH and SKF enacted the workshop. KM, YA and JDB analyzed the survey results. RK, SKF, KM, JDB, GG and CAH wrote the manuscript. All authors read and approved the final manuscript.

### Funding

This research was supported by a Small Business Innovation Research Grant #2R44NS108850-03A1 from the National Institute of Neurological Disorders and Stroke of the National Institutes of Health. CAH and GG are Principal Investigators on this grant.

### Competing interests

GG is a co-founder and co-owner of Backyard Brains, Inc., a company that manufactures and sells neurorobots like those described in this manuscript. CAH, MG, AH, and GG are employed by Backyard Brains, Inc. The remaining authors declare that the research was conducted in the absence of any commercial or financial relationships that could be construed as a potential conflict of interest.

## Supplemental Material

Supplemental Material 1

Lesson 1 - Teacher Guide and Student Handouts

Supplemental Material 2

Lesson 2 - Teacher Guide and Student Handouts

Supplemental Material 3

Lesson 3 - Teacher Guide and Student Handouts

Supplemental Material 4

SpikerBot Primer for Teachers

Supplemental Material 5

Pre/Post Survey

