## Supplementary material for "Building Brains for Robots: A Hands-On Approach to Learning Neuroscience in the Classroom": Lesson 1

TEACHING GUIDE

### Lesson 1: Neurons and Synapses

#### Lesson Goals

- Create a brain that can respond to changes in the environment
- Create a brain that can follow a moving target
- Create a brain that can explore on its own

#### Neuroscience Concepts

- Brains are made of **neurons**
- Neurons are connected by **axons** and **synapses**
- Neurons communicate with electrical signals called **action potentials** or **spikes**

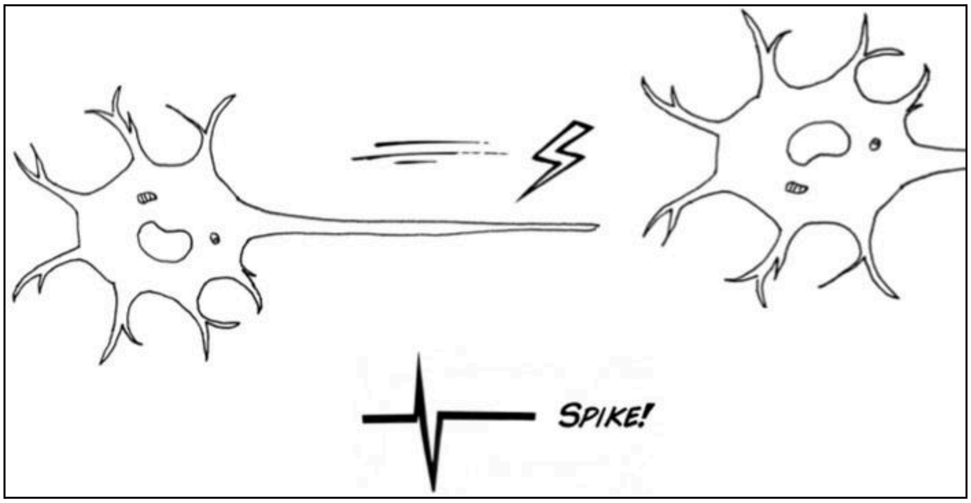


- Sensory neurons respond to **sensors** like eyes, ears and whiskers (or cameras, microphones, and distance sensors, in a robot)
- Motor neurons activate **effectors** like muscles and glands (motors, speakers, LEDs)
- Different **types** of neurons have different patterns of activity, eg. **quiet** and **bursting**

#### Why It Matters

Neurons and synapses are the fundamental building blocks of neural networks (brains and AIs). The function (behavior) of a neural network depends entirely on its structure (connectome), i.e. the types of neurons present and the way they are connected. This lesson teaches students how to build neural networks that make the SpikerBot produce a range of fun, life-like, sensory-guided behaviors.

#### Brain Design Basics Demonstration

See the [SpikerBot Primer] to learn how to install the app, connect to the SpikerBot, and navigate the app to create neural networks. We recommend teachers install the SpikerBot app, connect the computer to the SpikerBot's WiFi, and have the app open on student laptops before beginning lessons. Alternatively, teachers may opt to teach their students how to set up the SpikerBot app and connect to the SpikerBot as well.

After sharing background information on neurons and synapses, demonstrate for the students how to use the SpikerBot app to build a neural network. By the end of the demo, students should understand how to:

- **Switch between the Main Menu, Runtime mode, and Design mode**
  - Main Menu to Runtime mode: Create a new brain or select an existing brain. Click the 'Runtime' button on the bottom left of the screen. The 'Runtime' button will turn green before entering Runtime mode. Be patient.
  - Runtime mode to Design mode: Click on the 'Design' button on the bottom left of the screen.
  - Design mode to Runtime mode: Click 'Save' to save changes to brain design, then 'Runtime' on the bottom left of the screen.
  - Runtime mode to Main Menu: Click 'Menu' on bottom right of screen.

- **Create, move, or delete a neuron in Design mode**
  - Create a neuron: click anywhere in the brain and select the type of neuron.
  - Move a neuron: click on the neuron, then 'Move', then a new location in the brain.
  - Delete a neuron: click on neuron, then 'Delete'.

- **Create a synapse, modify synaptic weight, or delete a synapse**
  - Create a synapse and modify synaptic weight
    1. Sensor to neuron: click on the orange square next to a sensor (eyes, mic, or whiskers), click on neuron, select visual/distance/frequency preference depending on the input sensor, then 'Confirm'.
    2. Neuron to neuron: click on the first neuron, then 'Synapse', then the second neuron. Choose excitatory or inhibitory. Change the synaptic weight if desired, then 'Confirm'.
    3. Neuron to effector: click on neuron, then 'Synapse', then orange square next to the effector (motor/wheel or speaker). Adjust the synaptic properties if desired, then create synapse (motor or sound output synapse depending on the effector).
  - Delete axon:
    1. Sensor to neuron OR neuron to effector: repeat the same process as to create an axon, but in the final step click 'Delete synapse', then 'Confirm'
    2. Neuron to neuron: repeat the same process as to create an axon, then change synaptic weight to 0, then 'Confirm'.

#### Brain Design Challenges

After the demo, students can independently approach the Brain Design Challenges below, using the Lesson 1 Student Handout. There are many solutions to each Challenge, because many neural networks can perform the required behaviors in different ways. Example solutions are provided below each challenge.

##### Challenge #1: Design a brain that can avoid obstacles

Solution: A single quiet neuron (1) detects nearby objects and drives the motors backward. This simple circuit will prevent the robot from crashing into obstacles like walls.

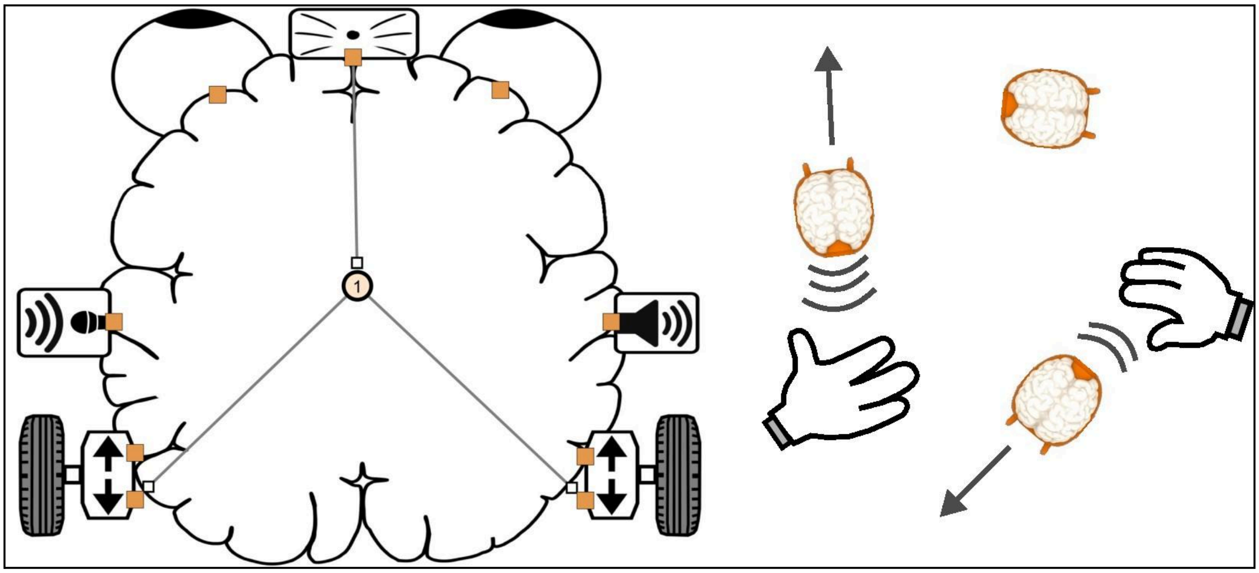


***Lesson 1 Exercise 1 (L1E1).*** *Obstacle avoidance.*

Design: Create a new brain. Click anywhere in the brain to create a quiet neuron. Click on the orange square next to the distance sensor (whiskers), then click on by the neuron. Select 'medium' distance preference, then 'Confirm'.

Create synapses between the neuron and the backward-going motors on both sides of the robot. Click on the neuron and select 'Synapse'. Next, create the synapse by clicking on the orange square next to the backward facing arrow of the left motor. Choose a synapse with a weight (strength) of 50, then 'Confirm'. Repeat this process on the right side.

Click 'Save', then 'Runtime' to see the robot operate based on this brain design.

**Challenge #2: Design a brain that can follow a moving target**

Solution: Two quiet neurons (1-2) detect red on the left or right. Two quiet neurons (3-4) drive the left or right motor forward. Red-detection on the left (1) drives the right-side motor (4) forward, and vice versa. Together, the four neurons form a small neural network that will turn the robot toward red objects in its peripheral vision, and approach them if they're straight ahead. This is an important example of how the structure of a biological system enables its function.

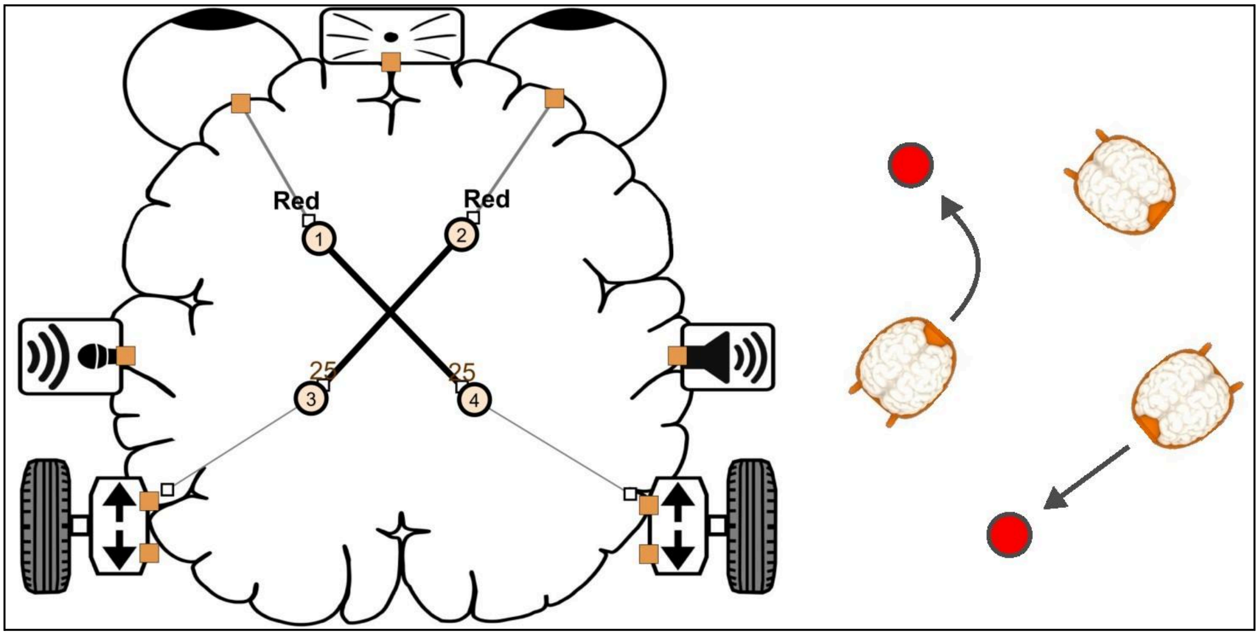


***Lesson 1 Exercise 2 (L1E2).*** *Target tracking.*

Design: In a new brain, create four quiet neurons. Neuron 1 will respond to the color red from the left eye (the left side of incoming camera frames) while neuron 2 will respond to red from the right eye. Click on the orange square next to the left eye followed by neuron 1. Select 'red' as the visual preference. This will create a synapse between the left eye and the neuron, with the word 'Red' written on it. Repeat this process to extend an axon from the right eye to neuron 2.

Neuron 1 will stimulate Neuron 4, which in turn will drive the right motor forward. To set this up, click on neuron 1, select 'Synapse', then click neuron 4. Choose an 'Excitatory' synapse with a weight of 25. Next, extend a synapse from neuron 4 to the forward-going motor on the right side by clicking on neuron 4, then 'Synapse', then the orange square by the forward-facing arrow of the right motor. Choose a synaptic weight of 25.

Conversely, Neuron 2 will excite Neuron 3, which will drive the left motor forward. To set this up, click on neuron 2, select 'Synapse', then neuron 3. Choose an 'Excitatory' synapse with a weight of 25. Next, extend a synapse from neuron 3 to the forward-going motor on the left side by clicking on neuron 3, then 'Synapse', then the orange square by the forward-facing arrow of the left motor. Choose a synaptic weight of 25.

Click ‘Save’, then ‘Runtime’ to see the robot operate based on this brain design.

Troubleshooting: If the robot is having trouble following the object, try moving the object a bit slower so that the robot's camera has time to detect the object/color and respond. You can also change how strongly the motors are activated, to prevent the robot turning too fast and losing sight of its target.

#### Challenge #3: Design a brain that can move around on its own

Solution: Five burst-generating neurons (1-5) drive the motors forward or backward and combine in unpredictable combinations. This neural network moves the robot around randomly.

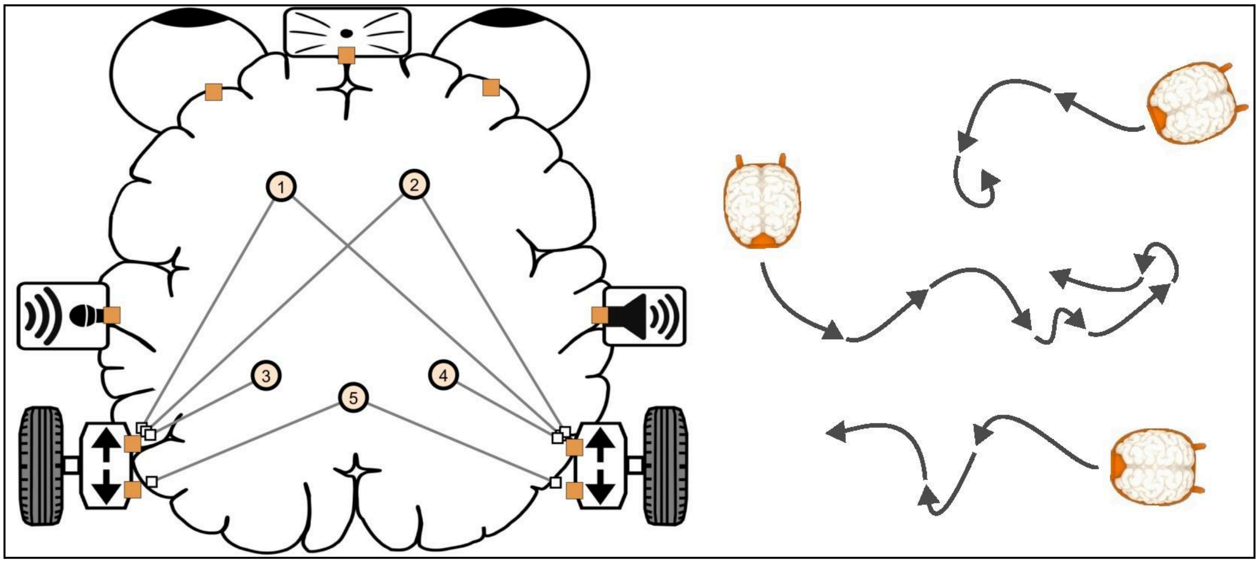


***Lesson 1 Exercise 3 (L1E3).*** *Exploration.*

Design: In a new brain, create several burst-generating neurons by clicking anywhere in the brain and selecting the neuron type ‘Generates bursts’ for each neuron. Randomly connect the bursting neurons to the motors by selecting a neuron, clicking ‘Synapse’, then the orange square next to any motor (front or back, and left or right).

Click ‘Save’, then ‘Runtime’ to see the robot operate based on this brain design.

Troubleshooting*:* To improve the robot’s exploration range, make the forward-going motor synapses stronger than the backward-going ones (e.g. 75 forward vs 25 backward). You can also add an obstacle avoiding neuron (L1C1) to prevent the robot getting stuck.

##### Challenge #4: Design a brain that blinks and makes sounds when it sees a cup

Solution: One quiet neuron (1) detects coffee cups. Four quiet neurons (2-5) produce different tones and colors. The cup detecting neuron activates the tone-generating neurons.

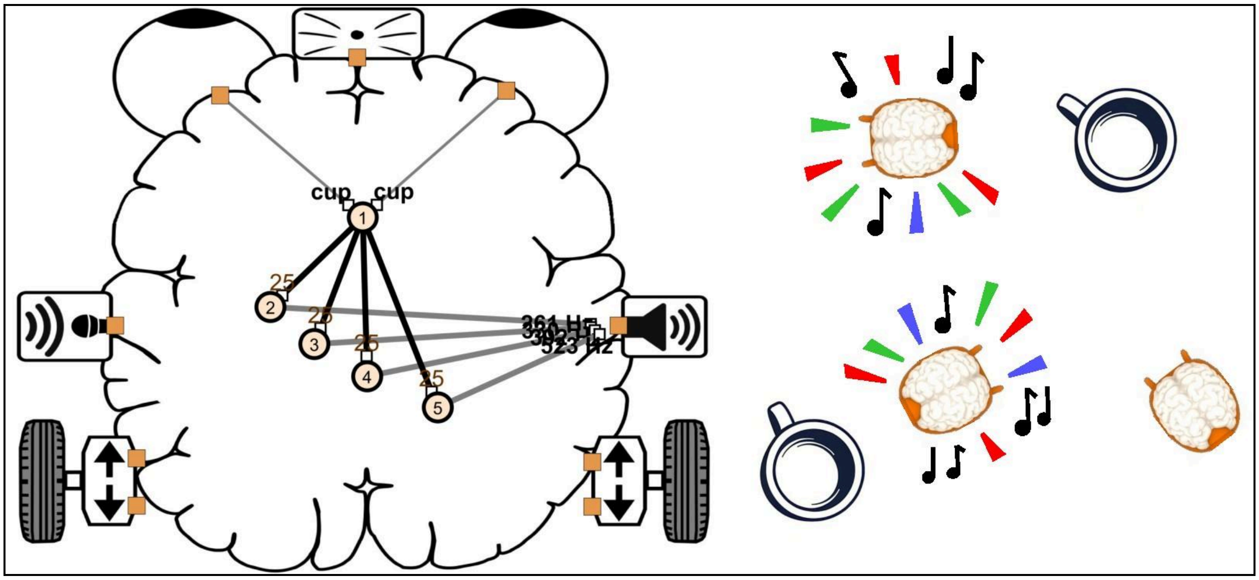


***Lesson 1 Exercise 4 (L1E4).*** *Blink and beep at cups.*

Design: In the Main Menu, create a new brain, then select ‘GoogLeNet’ from Trained Networks before you click ‘Runtime’. GoogLeNet is a neural network trained to recognize objects (see Primer). Go to Design. Add a quiet neuron. Click on the orange square next to either eye, and then on the new neuron. Select ‘cup’ as the visual preference. Repeat for the other eye.

Add 4 additional neurons. These will activate the speaker and blink the lights. Connect each neuron to the speaker by selecting the neuron, then ‘Synapse’, and then the orange square next to the speaker. Select output frequency (human speech is 50-500 Hz). To also connect the neuron to the robot’s lights, select the neuron, click ‘Specials’ and choose which color to blink.

Finally, connect the cup detector neuron (1) to the sound and light output neurons (2-5). Select the cup detector neuron, click ‘Synapse’, then click on an output neuron. Select ‘Excitatory’ to set the weight (strength) of the synapse to 25 - this is just under the threshold to reliably trigger an action potential, and will make the output more variable. If the weight of the synapse is higher, the output neurons will be maximally activated by the cup neuron and will always produce the same tone output, which students find frustrating.

Troubleshooting: Make sure your robot recognizes the cup. Try holding the cup in front of the robot and see if it responds with sounds and lights. Observe the cup from the robot’s perspective (using the camera images displayed in the app) to make sure the cup is fully visible. If the brain is having trouble detecting the cup, try different cups and improve the room lighting.

#### Resources

- Lesson 1: Video

#### Reading

- Harris et al. (2020) Neurorobotics Workshop
- Valentino Braitenberg (1984) Vehicles
- Jeffrey Krichmar (2018) Neurorobotics

#### Standards Alignment

- **NGSS LS1.A: Structure and Function.** The function of a neural network depends on its structure. For example, visually guided approach behavior (function) derives from crossing streams of information from the left side of the visual field to the right side of the body, and vice versa (L1E2).
- **NGSS ETS1.B: Developing Solutions to Real-World Problems.** Accurate color and object processing in video is challenging, especially with a moving camera. Learn how to modify the brain or the environment to better support visually guided approach behavior.

STUDENT HANDOUT

### Lesson 1: Neurons and Synapses

#### Lesson Goals

- Create a brain that can respond to changes in the environment
- Create a brain that can follow a moving target
- Create a brain that can explore on its own

#### Neuroscience Concepts

- Brains are made of **neurons**
- Neurons are connected by **axons** and **synapses**
- Neurons communicate with electrical signals called **action potentials** or **spikes**

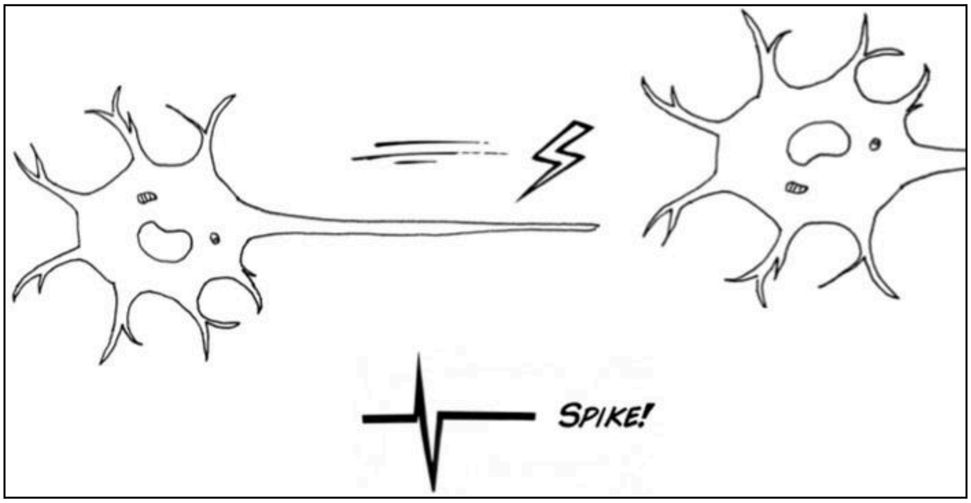


- Sensory neurons respond to **sensors** like eyes, ears and whiskers (or cameras, microphones, and distance sensors, in a robot)
- Motor neurons activate **effectors** like muscles and glands (motors, speakers, LEDs)
- Different **types** of neurons have different patterns of activity, eg. **quiet** or **bursting**

**Lesson 1**

### Brain Design Exercises

##### Exercise #1: Design a brain that avoids obstacles

Hint: Add a neuron that is activated by the distance sensor and moves the robot backward.

##### Exercise #2: Design a brain that can follow a moving target

Hint: Let the left eye activate motors on the right side and vice versa.

##### Exercise #3: Design a brain that can move on its own

Hint: Add neurons that generate bursts. Connect them to the motors. Observe what happens. Add the obstacle-avoiding neuron from Exercise #1 to avoid collisions.

##### Exercise #4: Design a brain that blinks and makes sounds when it sees a cup

Hint: Create a neuron that responds to cups (to do this, make sure the trained network GoogLeNet is selected in the Main Menu before you start). Add other neurons that produce sounds and lights. Connect the cup detecting neuron to the sound/light output neurons.
