## Supplementary material for "Building Brains for Robots: A Hands-On Approach to Learning Neuroscience in the Classroom": Lesson 2

TEACHING GUIDE

### Lesson 2: Decision Making

#### Lesson Goals

- Import previously designed neural networks into a new brain
- Configure a brain so it can choose effectively between different neural networks
- Evaluate a brain's ability to find a hidden object

#### Neuroscience Concepts

- Complex behaviors can be decomposed into simpler behaviors controlled by distinct neural networks
- A brain structure called the **basal ganglia** can **disinhibit** (activate) neural networks, one at a time, allowing the brain to choose what to do at any given moment (action selection)
- Inputs to the basal ganglia's input neurons ('**striatal neurons**') influence when, how often and for how long a particular neural network is selected

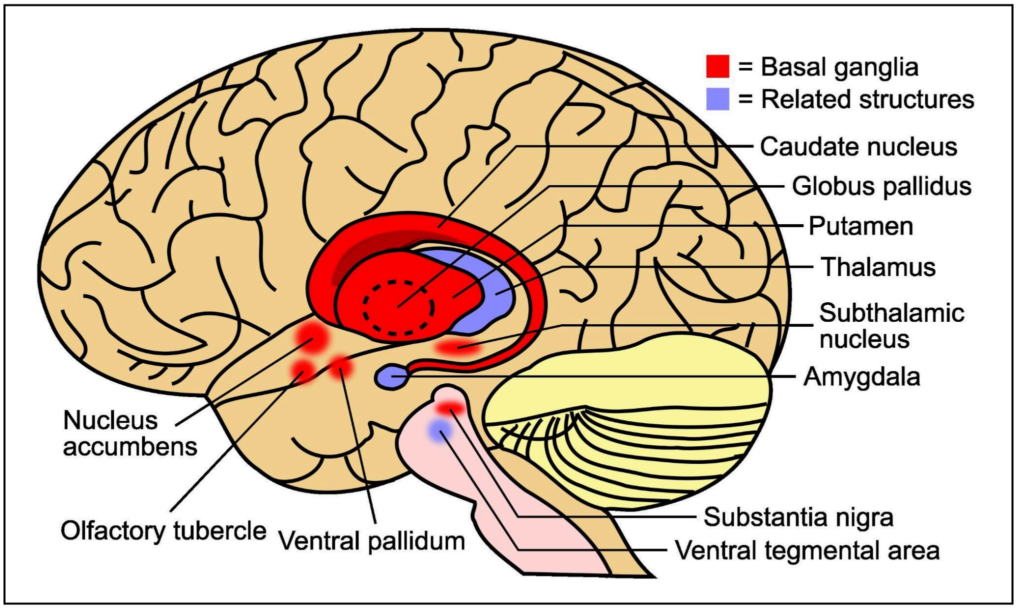


*The basal ganglia is a group of interconnected structures located deep within the brain. Its function is to provide selective inhibitory control over a wide range of neural networks.*

#### Why It Matters

The basal ganglia play a crucial role in controlling brain activity and motor output. Dysfunction of the basal ganglia is implicated in many brain disorders including anxiety, schizophrenia, Parkinson's disease and addiction. This lesson teaches students how to import neural networks (from the previous lesson) into a single brain and configure the brain's basal ganglia to select intelligently between them.

#### Basal Ganglia Basics

The basal ganglia uses strong, precisely targeted synaptic inhibition to keep neural networks throughout the brain quiet and under control. It can selectively release individual networks from this inhibition, allowing them to generate action potentials, respond to stimuli, and produce behavior. This is how vertebrate animals make choices.

In the SpikerBot app, all neurons can be assigned a unique basal ganglia network ID (A, B, C etc). The different networks are engaged in a **continuous competition to be selected** by the basal ganglia. Each network is associated with a level of **motivation** (selection likelihood) shown in the bottom right bar plot in Runtime mode. The network with the highest motivation is the currently selected network. Only one network can be selected at a time. When selection switches to a new network, this is indicated by a jump in its motivation, the network ID tag shown above the neurons in the network switching from black to white, and the thinning of previously thick gray lines connecting all neurons in the network.

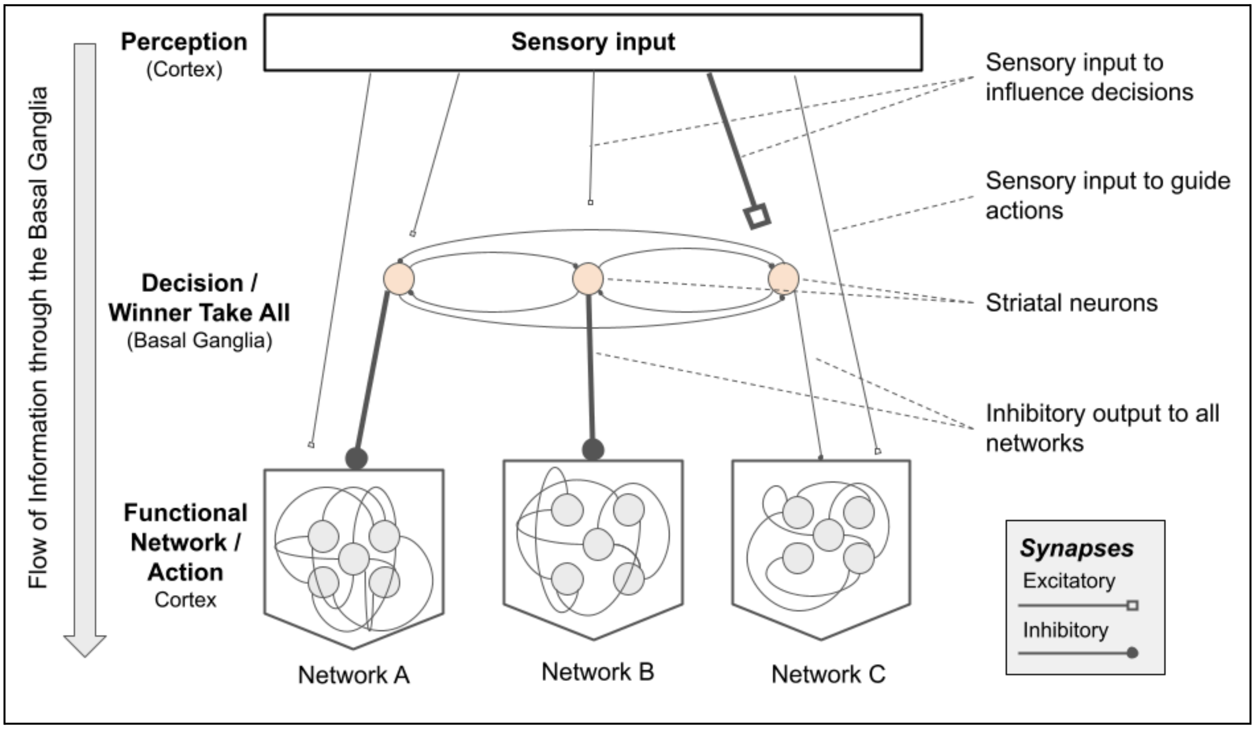


*Basal ganglia selecting between 3 networks. Sensory input excites the striatal neuron controlling network C, causing it to become selected (disinhibited). Thick axons indicate high activity.*

In the absence of input, the basal ganglia selects randomly between the available neural networks. However, the brain can influence the decision-making process by sending synaptic inputs to the basal ganglia's input neurons, which are called **striatal neurons**. Each striatal neuron controls a specific network ID. Any synaptic input to a striatal neuron, positive or negative, is applied to the network's motivation. By connecting the right sensory neurons to the right striatal neurons, the brain can ensure that the right network will be selected at the right time. For example, a red traffic light inhibits driving, whereas a green light stimulates driving.

#### Pre-Lesson Material Preparation

The brains in this lesson are designed to explore. Therefore, the robot needs to be in an arena to prevent it getting lost. To make an arena, cut strips of cardboard 5 cm tall and about 1 m long. Tape them together to form a square.

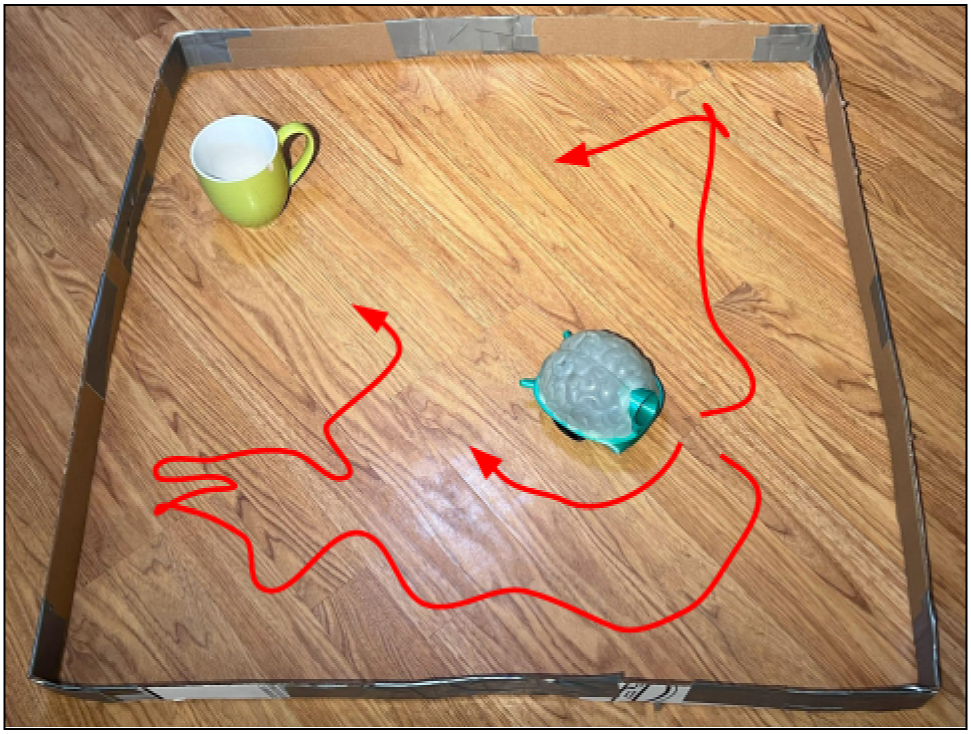

*The robot starts facing away from the target. The task is to find the cup.*

Make sure each student group has a timer and is able to use it. If students are working in groups of 3 or more, assign them roles e.g. materials (manage the robot and the arena), navigator (manage the laptop and app), and timer (manage the timer and log the times). If possible, one student can draw the path the robot takes in the arena. This will allow them to remember and compare the behaviors of different brains.

#### Brain Design Exercises

*See the [SpikerBot Primer] to learn how to install the app, connect to the SpikerBot, and navigate the app to create neural networks.*

The overall aim in this lesson is to build a brain that can find a cup. In Exercises 1-2, the brain will perform the task poorly, but by Exercises 3-4, it will find the cup in about a minute. We recommend Exercises 1-2 are performed together as a class. They can then independently approach Exercises 3-4, using the Lesson 2 Student Handout. Solutions and explanations are provided with each exercise.

Troubleshooting: If you return to the Main Menu after creating any of the following brains, you will need to select the Trained Network 'GoogLeNet' before reopening the brain. Without doing so, the Runtime button will turn red, preventing access to Runtime mode.

#### Exercise #1: Create a Brain that can Select Between Two Neural Networks

Step 1: Create a new brain with an obstacle-avoiding neuron (see Lesson 1, Exercise 1). The neuron will not be under the basal ganglia's inhibitory control and will always be ready to move the robot backward if an obstacle is detected.

Step 2: Click 'Import Brain', then click in an empty part of the brain. Select the spontaneously moving brain from Lesson 1 Exercise 3 (L1E3) from the dropdown menu and click 'Confirm'. The imported network is automatically assigned a network ID (A).

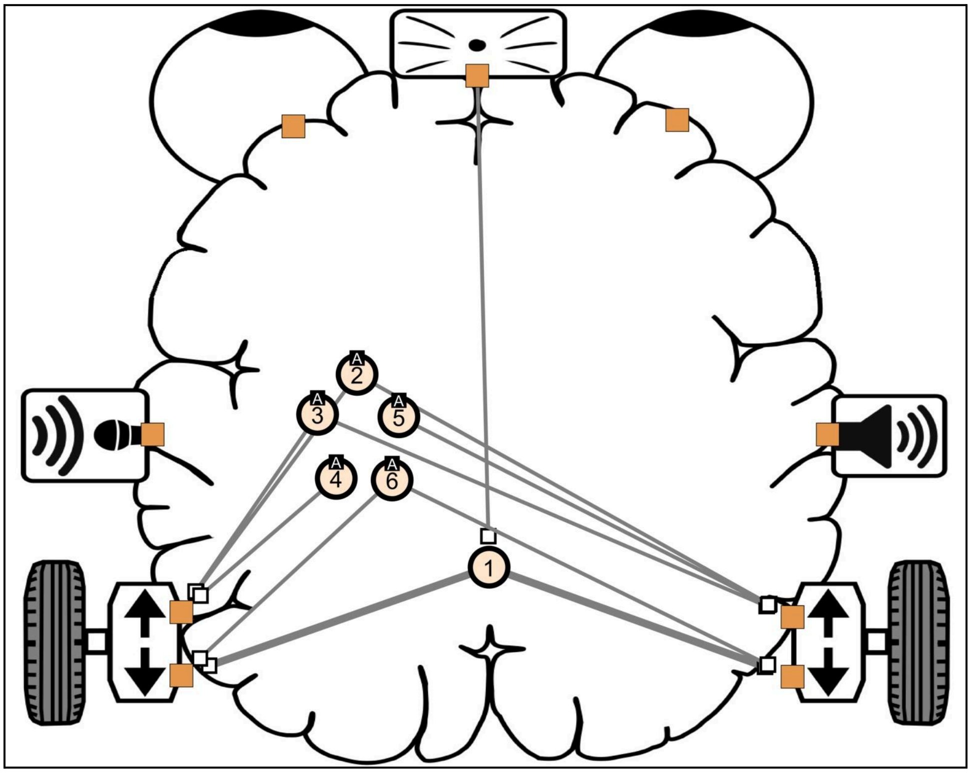


***Lesson 2 Exercise 1 (L2E1).*** *Step 2.*

Step 3: Next, click ‘Import Brain’ again, and click in another empty part of the brain. Select the cup-detecting brain from Lesson 1 Exercise 4 (L1E4), and click ‘Confirm’. The imported network is automatically assigned a network ID (B).

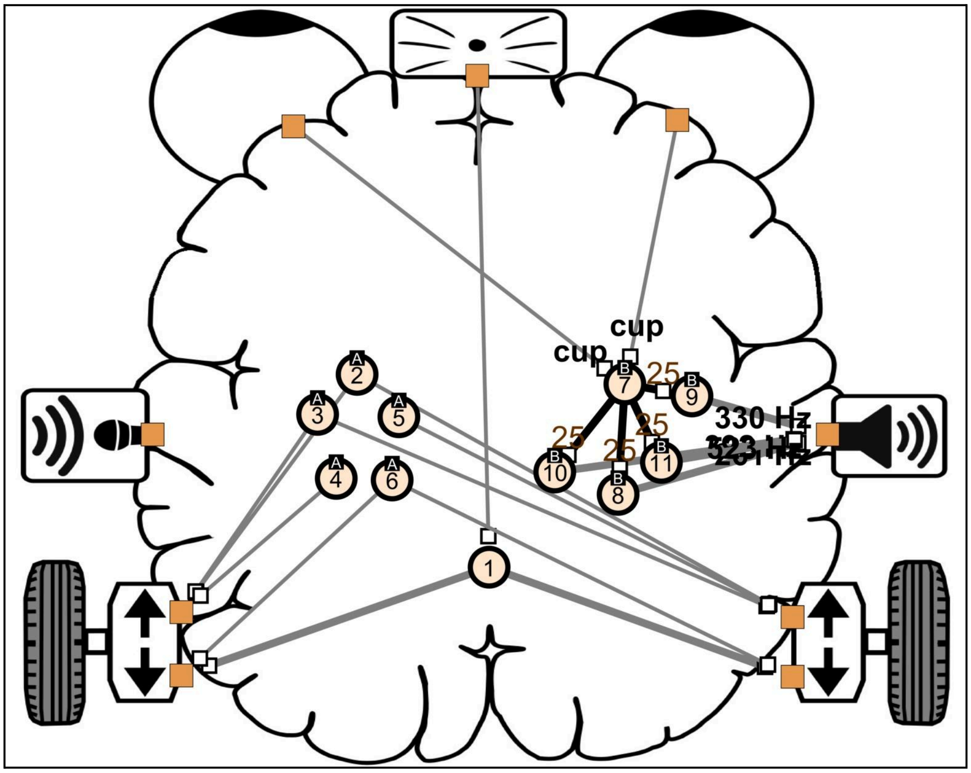


***Lesson 2 Exercise 1 (L2E1).*** *Step 3.*

This will produce a brain with two neural networks, each producing a different behavior. Network A will produce **exploratory** behavior when selected, while Network B will **indicate** whether a cup has been found with sounds and lights.

<u>Step 4:</u> Instruct students to place the robot in the arena facing away from the cup and go to Runtime mode to observe its behavior. When network A is selected the right-side bar plot shows high motivation for A (purple bar) and the neurons in network A are active (blinking). When network B is selected the orange bar is high and the neurons under B will be active (blinking) **if** a cup is in view. If a cup is not in view when network B is selected, the robot will simply stay still.

The aim is for students to see that without synaptic input, the basal ganglia will select each network ~50% of the time whether or not the cup is in view. It will therefore be slow to explore, and will rarely stop to indicate when it sees the cup.

#### Exercise #2: Modify the Brain so it Stops and Indicates when it Sees a Cup

To make the brain switch to network B (cup-indication) only when a cup is actually in view, it is necessary to add 3 more neurons: two striatal neurons - one for each imported network - as well as a cup-detecting neuron. The cup-detecting neuron needs to inhibit the striatal neuron controlling network A (exploration) and excite the striatal neuron controlling network B (cup-indication).

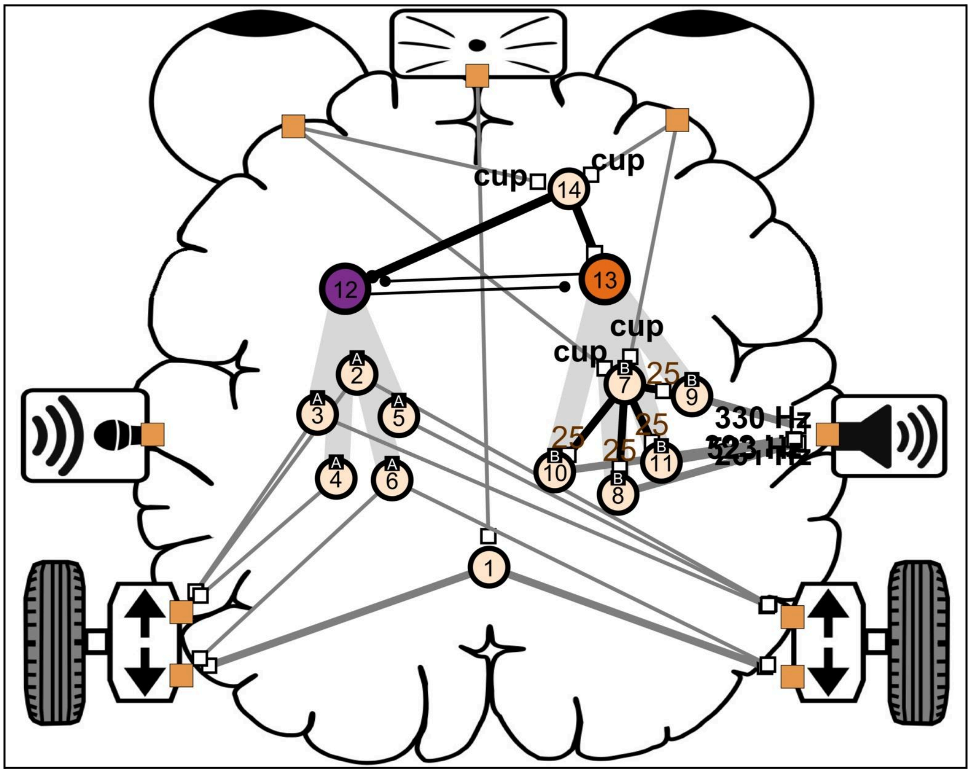


***Lesson 2 Exercise 2 (L2E2)***

<u>Step 1:</u> Add a striatal neuron (12) above the neurons in network A. Before clicking ‘Confirm’ to create the striatal neuron, ensure that ‘A’ is selected as the ‘id’ in the dropdown menu. The striatal neuron will appear purple. A gray cone will connect this striatal neuron to the network A neurons.

<u>Step 2:</u> Next, add a second striatal neuron (13) above the neurons in network B. Before clicking ‘Confirm’ to create the striatal neuron, ensure the ‘B’ is selected as the ‘id’ in the dropdown menu. This striatal neuron will appear orange. A gray cone will connect this striatal neuron to the network B neurons.

<u>Step 3:</u> To set up the cup-detecting neuron, create a quiet neuron (14). Click on the orange square next to the left eye followed by this quiet neuron. Select ‘cup’ as the visual preference. A synapse should be extended from the left eye to the neuron, with the word ‘cup’ written on the synapse. Repeat this process to create a synapse from the right eye to the same neuron (14).

Step 4: To make the cup-detecting neuron **excite** the striatal neuron controlling network B (indication behavior), click on neuron 14, select 'Synapse', then select the striatal neuron controlling network B (neuron 13) to extend an axon from neuron 14 to neuron 13. Choose an 'Excitatory' synapse with weight of 25. A line connecting neuron 14 to neuron 13, ending in a white square, will appear.

Step 5: To make the cup-detecting neuron **inhibit** the striatal neuron controlling network A (exploration behavior), click on neuron 14, select 'Synapse', then the striatal neuron controlling network A (neuron 12) to extend an axon from neuron 14 to neuron 12. Choose an 'Inhibitory' synapse with weight of -25. An axon from neuron 14 to neuron 12, ending in a black circle, will appear.

Step 6: Instruct students to place the brain in the arena, facing away from the cup, and observe its behavior. The aim is for students to see that the robot will now stop moving when it sees the cup and switch to its indication behavior. It will not begin moving again as long as the cup is in view since the exploration network is being inhibited. However, the indication network (B) will still be selected ~50% of the time when the cup is not in view, which (in the absence of a cup) leaves the robot motionless for ~20 s, and increases the time it takes the brain to find the cup.

#### Exercise #3: Modify the Brain So It Explores in the Absence of Cups

To make the brain choose exploration whenever the cup is not in view, it is necessary to add one more neuron: a spontaneously active neuron that stimulates network A (exploration) and inhibits network B (indication).

Step 1: To set up this biasing neuron, create a 'Highly Active' neuron in the top left corner of the brain (neuron 15).

Step 2: To make the biasing neuron **excite** the striatal neuron controlling network A (exploration), click on neuron 15, select 'Synapse', then the striatal neuron controlling network A (12) to extend an axon from neuron 15 to neuron 12. Choose an 'Excitatory' synapse with a weight of 25. A line connecting neuron 15 to neuron 12, ending in a white square, will appear.

Step 3: To make the biasing neuron **inhibit** the striatal neuron controlling network B (indication), click on neuron 15, select 'Synapse', then the striatal neuron controlling network B (13) to extend an axon from neuron 15 to neuron 13. Choose an 'Inhibitory' synapse with a weight of -25. A line connecting neuron 15 to neuron 13, ending in a black circle, will appear.

Step 4: Instruct students to place the robot in the arena, facing away from the cup, and observe its behavior. The brain will now choose to explore whenever the cup is not in view.

<u>Step 5:</u> Students should now time how long it takes the brain to find the cup. Run 5 trials to get an average time. The brain should be able to find a cup in a 1x1 m arena in 1-2 min.

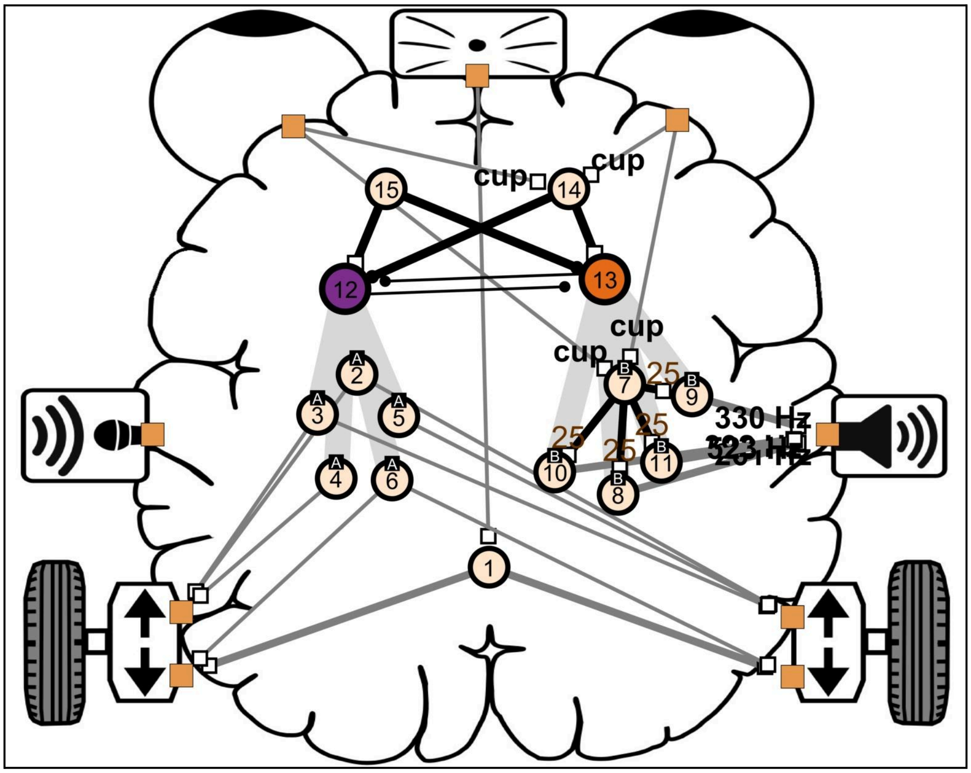


***Lesson 2 Exercise 3 (L2E3)***

*<u>Troubleshooting</u>: Make sure the robot can see the cup from different angles. Use more than one cup if your robot is having trouble detecting it.*

#### Exercise #4: Modify the Brain To Improve its Performance

There are many things students can do to improve their brain's performance:

- Increase or decrease the number of neurons in the exploration network (A).
- Change the rate and duration of bursting in network A by modifying the neurons' b and c properties, respectively.
- Increase or decrease the strength of the motor output synapses.
- Change the strength of the synapses that excite or inhibit the striatal neurons (e.g. increase input to the striatal network B neuron to make the brain switch to indication more rapidly).

#### Why is it called 'basal ganglia'?

The basal ganglia consists of a large number of mostly inhibitory neurons organized into groups or 'ganglia'. The striatum is the first and largest ganglion. It's the 'input ganglion', but it's also the ganglion in which the decision-making happens. Neurons in the striatum inhibit each other, creating a 'winner-take-all' dynamic, where the winning grouping of neurons decide what to do (which neural network gets released from inhibition) at any given moment. This action selection process is profoundly influenced by inputs from the rest of the brain.

#### Resources

- 2-Minute Neuroscience: Basal Ganglia

#### Reading

- Grillner et al. (2007) Neural Bases of Goal-Directed Locomotion in Vertebrate
- Prescott et al. (2006) A Robot Model of the Basal Ganglia: Behavior and Intrinsic Processing
- Bolado-Gomez and Gurney (2013) A Biologically Plausible Embodied Model of Action Discovery

#### Standards Alignment

- **NGSS LS1.A: Structure and Function.** Inhibitory control that can be selectively removed from specific networks (structure) enables decision-making (function).
- **NGSS ETS1.B: Developing Solutions to Real-world problems.** Modifying brains and environments to improve poor decision-making.
- **Computational Thinking (CT).** Breaking a complex behavior down into components.

**STUDENT HANDOUT**

### Lesson 2: Action Selection

The **basal ganglia** is a brain structure that enables us to make decisions. It does this by 'disinhibiting' (activating) neural networks one at a time.

Inputs to the basal ganglia's '**striatal neurons**' strongly influence when and for how long a particular neural network is selected.

In the SpikerBot app, neurons are assigned a basal ganglia ID. Only one ID is selected (chosen) at any given time, and neurons can only fire spikes if their network ID is currently selected.

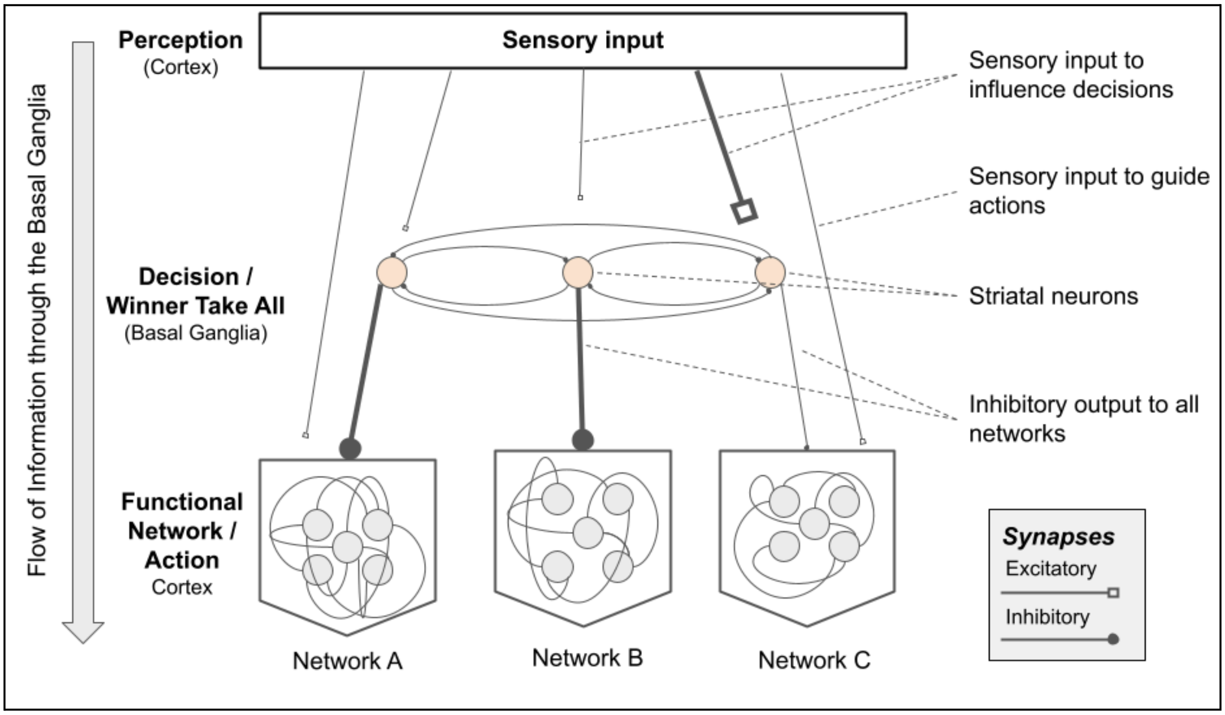


*Schematic diagram of a brain that can select between 3 networks. Network C is selected.*

#### Lesson 2
### Brain Design Exercises

The overall aim is to build a brain that can find a cup. In Exercise 1-2 it won't work so well, but by Exercise 3, your robot should be able to find a cup in 1-2 minutes.

##### Exercise #1: Combine Networks with Different Behaviors into a Single Brain

Create a brain with a single obstacle-avoiding neuron. Import two neural networks from the previous lesson: the randomly exploring L1E3 and the cup-crazy L1E4. Run the brain. What are your observations about how it moves? Is it able to find the cup in under 2 minutes?

##### Exercise #2: Modify the Brain so it Stops and Indicates when it Sees a Cup

The problem with the brain in the previous exercise is that its basal ganglia is not using any information to make decisions. Let's give it some synaptic input. Add two 'striatal' neurons (one for network A and one for network B) and a single cup-detecting neuron. Make the cup-detecting neuron inhibit the striatal network A neuron and excite the striatal network B neuron. Run the brain. What are your observations about how it moves? Is it able to find the cup in under 2 minutes? Did the striatal neurons make a difference?

##### Exercise #3: Modify the Brain So It Explores in the Absence of Cups

Use a highly active neuron to make the brain choose exploration (network A) in the absence of cups. Make the neuron excite the striatal network A neuron and inhibit the striatal network B neuron. What are your observations about how the robot moves? Can it find the cup in under 2 minutes? Did adding the internal (biasing) neuron make a difference?
