## Supplementary material for "Building Brains for Robots: A Hands-On Approach to Learning Neuroscience in the Classroom": Lesson 3

TEACHING GUIDE

### Lesson 3: Learning

#### Lesson Goals

- Evaluate a neural network that can recognize visual features in a new environment
- Train a neural network to navigate to specific locations in a new environment

#### Neuroscience Concepts

- Neural networks **learn** by changing the strength (weight) of synaptic connections between neurons
- **Associative learning** is when the brain learns to identify patterns in data without being explicitly told what the patterns are
- **Reinforcement learning** is when the brain learns to perform actions that lead to desired outcomes and avoid actions that lead to undesired outcomes

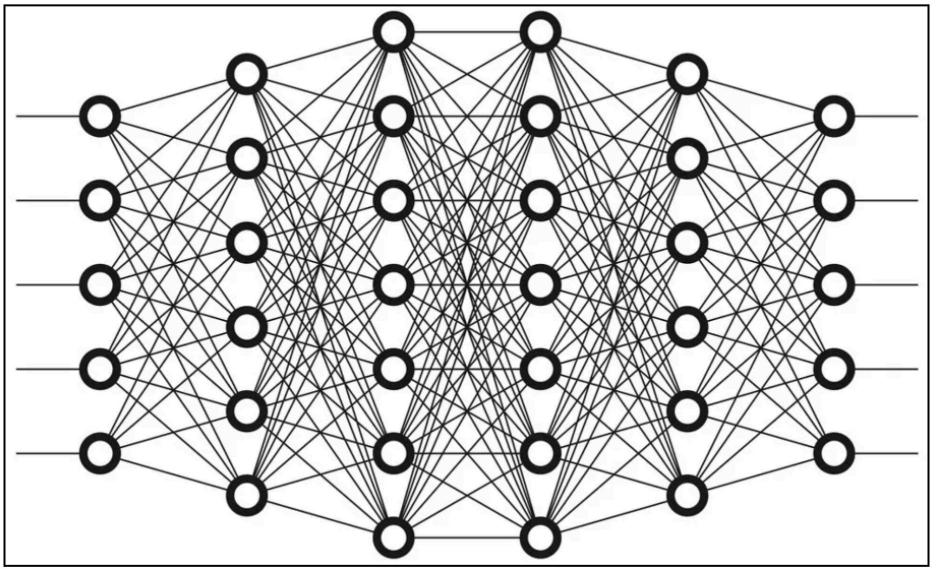

*A deep neural network has multiple layers of neurons. The neurons in one layer are connected to the neurons in the next layer, and so on. The network learns by adjusting the strengths of its synaptic connections to minimize errors.*

#### Why It Matters

The ability of neural networks to learn is what makes them so powerful and interesting. Brains can learn to **identify patterns** and to **achieve goals**. The following exercises teach students how to work with one neural network that can recognize locations in image data, and train another network to navigate the robot to a specific location. Students will gain a practical understanding of how neural networks can be trained to behave intelligently.

#### Deep Learning Basics

In this lesson, we will use **deep learning**, a technique for training neural networks with many layers of neurons (hence ‘deep’). While there are differences between deep learning and the learning that goes on in brains, the outcomes are very similar: neural networks learn to **recognize patterns** and **make decisions**.

**Classifier networks** (‘associative’ or ‘unsupervised’ learning networks) are used to identify recurring patterns in complex data, such as the SpikerBot’s video feed. The object-classifying network GoogLeNet, which we used in the previous lessons, is a classifier network. Classifier networks can be used to find patterns in all kinds and combinations of data, and have numerous applications in audio and video technology, finance, medicine, etc.

In this lesson, we will train a classifier network from scratch. First, camera images will be recorded and grouped together based on how similar they are. This “clustering” process usually identifies about 20 recurring places or viewpoints (e.g. specific pieces of furniture observed from particular angles). Then, deep learning will be used to train a classifier network to classify new camera images from the same environment into one of those 20 place categories. This creates 20 ‘place cells’ that indicate where the robot is located and which way it’s facing.

**Agent networks** (or simply ‘agents’) are neural networks that learn how to act to obtain rewards. For example, the AI that first mastered Atari video games in 2015 was a ‘deep Q’ agent network. Agent networks can be trained on any data where the current state (e.g. location) depends causally on the previous action (e.g. movement). Agent networks have applications in gaming, social media, home automation, robotics, and more.

Agent networks take an input representing the current state (e.g. location) and produce an action (e.g. leftward turn). They are trained by first creating a model of the environment called a Markov Decision Process (MDP). The MDP describes how likely the robot is to transition from any one state to any other. Then, deep learning is used to discover action sequences that will take the robot to the goal state.

For this lesson, students will use a classifier network to estimate the robot’s current state (location and orientation). Then, deep learning will be used to discover motor output sequences that will drive the robot to a specific location in the arena.

**Naming and Loading Trained Networks.** Neural networks trained with deep learning are available under Trained Networks in the Main Menu, and must be selected **before** clicking ‘Runtime’ to enter Runtime mode. Classifier network names should indicate the environment they represent (e.g. ‘office’). Agent network names should indicate the goal state they will attempt to navigate to (e.g. ‘cups’). Agent names will be automatically prefixed with the name of the associated classifier network (e.g. ‘office---cups’).

#### Pre-Lesson Data Preparation

In this lesson, we will train a classifier network and an agent network to perform the task from the previous lesson: finding cups in a 1x1 m arena.

Deep learning requires a lot of data. For this lesson, students will use **camera images** and **motor commands** recorded by the SpikerBot as it explores the arena. This data can be recorded by students during Lesson 2 or separately by the teacher, but needs to be recorded and processed **before** Lesson 3. It is also important that the recording and testing environments are as similar as possible (try not to move the arena).

Use the autonomously exploring brain L1E3 to collect training data (adding an obstacle-avoiding neuron is recommended). To record data, select ‘Record Data’ under ‘App’ settings in the Main Menu. Make sure ‘Record Data’ is still highlighted when you press ‘Runtime’ as some actions in the Main Menu, such as creating a new brain, causes menu settings to return to default values. The SpikerBot saves all recorded data to a local folder: ‘Documents\MATLAB\Datasets’.

You need to record data for at least 30 minutes to have enough data for training. More data will increase the processing time but will also improve performance. Deep learning is highly scalable, meaning it continues to improve no matter how much training data it’s given.

After you have finished recording data you need to train a classifier network. Press ‘Menu’ to return from Runtime mode to the Main Menu. Then, press ‘Learning’ to enter the deep learning interface. Provide a name for your new network and press ‘Train classifier network’ to begin processing training data. Your trained classifier network will be available under ‘Trained Networks’ in the Main Menu.

The deep learning process requires around 2 hrs per 30 min of recorded data. If successful, it will output ‘Ready to train agent network’. At this point, the laptop can be handed over to students, who can then approach the exercises below independently. You are free to return to the Main Menu menu by clicking ‘Exit ML’ and turn the laptop off. You or your students can restore the processed training data by clicking ‘Load classifier network’ in the Learning interface.

#### Brain Design Exercises

The aim of this lesson is to have students train neural networks to more effectively perform the task from in the previous lesson - finding a cup in an arena. Use the same arena as in Lesson 2.

Share background information on deep learning and artificial intelligence. In particular, explain to students how the recorded training data has been pre-processed for them. Students can then approach the exercises below independently.

#### Exercise #1: Test the trained classifier network

In the Main Menu, create a new brain, select the trained classifier network under Trained Networks, then click Runtime, then Design. In Design mode, choose ‘Import Trained Network’ to import your classifier network. Test it by moving the robot around. Does it classify different locations consistently (do the same neuron respond)?

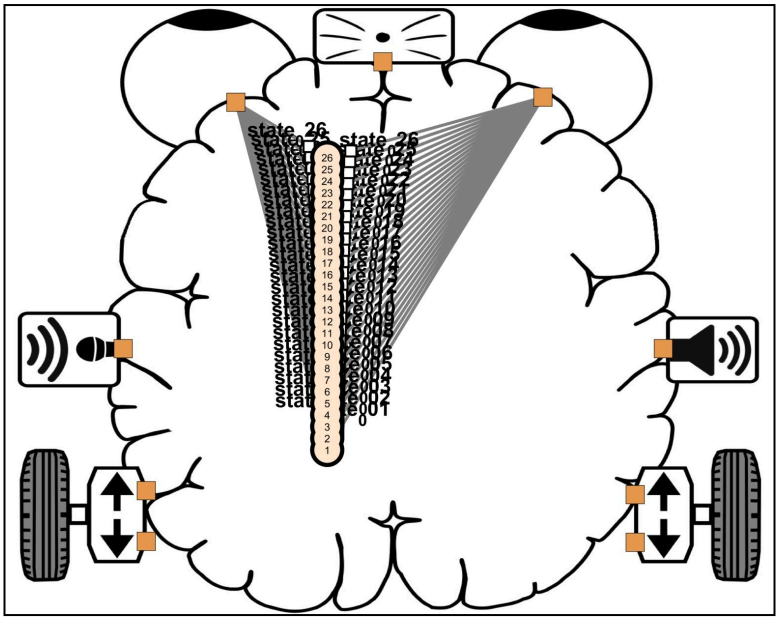

***Lesson 3 Exercise 1 (L1E3).*** *Classifier network imported into an empty brain. Only the output (classification) layer is shown.*

#### Exercise #2: Train an agent network to navigate to a specific location

In the Main Menu, press ‘Learning’. Select ‘Load classifier network’. A figure displaying a representative image from each identified location will be displayed.

The goal state is the location you want the robot to navigate to. To choose which states you want the robot to go to, use the figure showing representative images from each state or go to your Workspace folder (Documents\MATLAB\Workspace) to see all images. For this exercise, find states for which every image features a cup. Enter those states in the ‘Goal states’ input box, separated by spaces.

In the text input box that reads ‘Save agent network as’, enter a unique name for the network you are about to train. Then press ‘Train agent network’. This trains a neural network how to act in each state (location) in order to reach the goal state (cup).

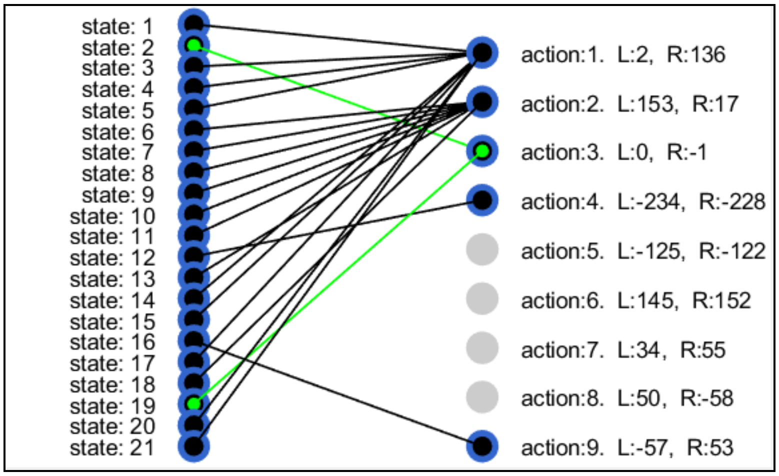


***Lesson 3 Exercise 2 (L3E2).*** *An agent network 'knows' which actions to take to reach a goal state (green dots and lines).*

Return to the Main Menu. Your trained networks should now be available under 'Trained Networks' (e.g. 'office---cups'). One way to visualize and test the trained network is to import it into an empty brain. To do this, create a new brain, go to Design mode, click 'Import Trained Network', click in the brain to place it, then select your trained network from the dropdown menu.

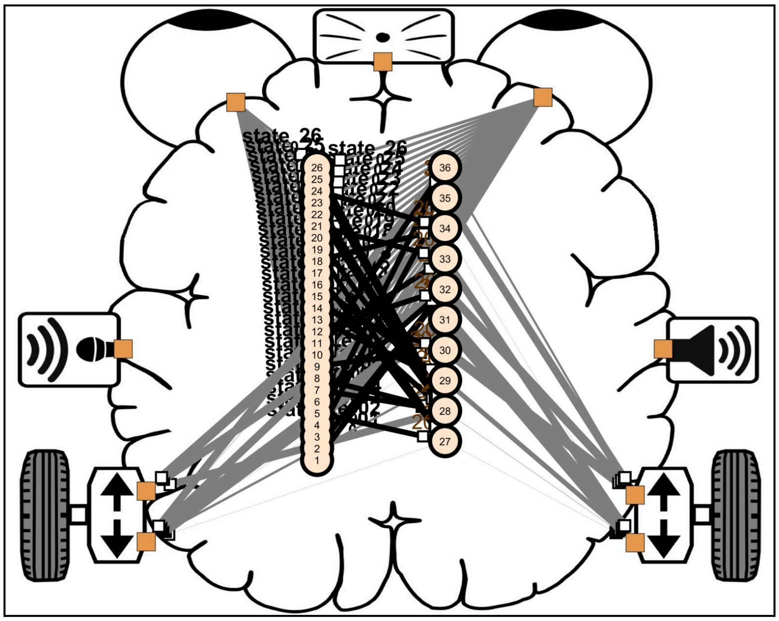

***Lesson 3 Exercise 2 (L3E2).***

How long does it take the trained net to find the cups? How does it compare to the L2E3 brain? We have found that L2E3 finds cups in about 54 s, whereas the trained agent network controlling L3E3 finds cups in about 27 s.

#### Standards Alignment

- *LS1.A:* Changing the synaptic weights of a neural networks (structure) leads to changes in neural network activity behavior (function)

**STUDENT HANDOUT**

### Lesson 3: Learning

Neural networks **learn** by changing the strength (weight) of synaptic connections between neurons. **Associative learning** is when the brain learns to identify patterns in data without being explicitly told what the patterns are (like learning to recognize a new face or place). **Reinforcement learning** is when the brain learns to perform actions that lead to desired outcomes and avoid actions that lead to undesired outcomes (like learning to ride a bike or play a game).

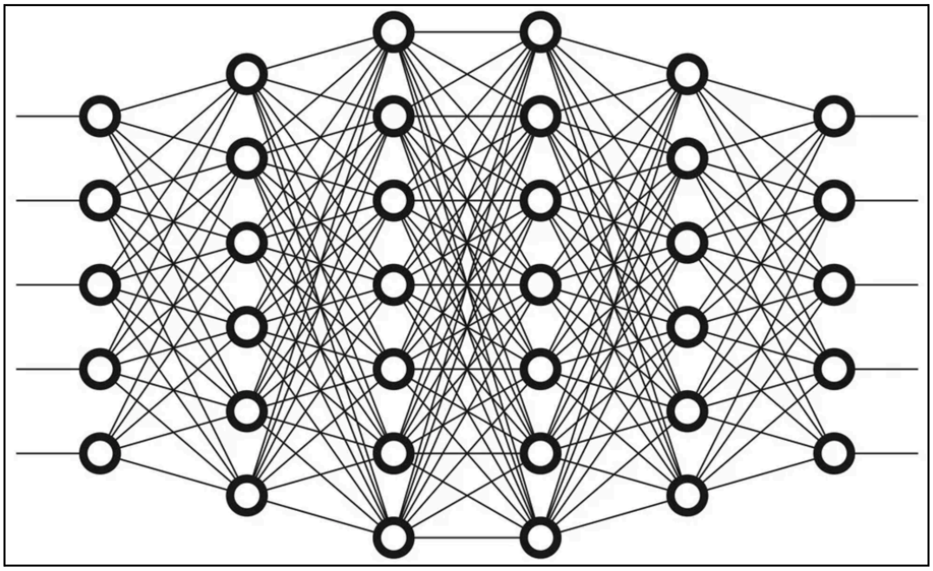

*A deep neural network has multiple layers of neurons. The neurons in one layer are connected to the neurons in the next layer, and so on. The network learns by adjusting the strengths of its synaptic connections to minimize errors.*

**Lesson 3**
### Brain Design Exercises

The overall aim here is to learn to load, train, test and work with large neural networks.

#### Exercise #1: Test the trained classifier network

Your teacher has trained a ‘classifier’ network to remember where it is. In this exercise you will import that network into an empty brain. Then you will move the robot around to see if the same neurons are consistently activated when the robot is in a certain location.

In the Main Menu, create a new brain, select the trained classifier network under Trained Networks, then click Runtime, then Design. In Design mode, choose ‘Import Trained Network’ to import your classifier network. Test it by moving the robot around. Does it classify different locations consistently (do the same neuron respond)?

#### Exercise #2: Train an agent network to navigate to a specific location

In this exercise, you will train an ‘agent’ (a network that maps locations to actions) that can navigate the robot to a specific location.

In the Main Menu, press ‘Learning’. Select ‘Load classifier network’. Enter goal states. Enter a unique name for the network. Press ‘Train agent network’ to train a network how to act in each state (location) in order to reach the goal state (cup).

When the training process is complete, return to the Main Menu and go to Design mode. Click ‘Import Trained Network’, click in the brain to select a location, then select your trained network from the dropdown menu to place it in the brain.

Test your brain, how is it performing?
