## Supplementary material for "Building Brains for Robots: A Hands-On Approach to Learning Neuroscience in the Classroom": Primer for Teachers

### SpikerBot Primer for Teachers

In today's world, it is essential that all students learn about **neural networks**:

- Our **brains** are **biological neural networks** that evolved to help us survive and thrive by shaping our **behavior**. To understand ourselves and treat brain disorders that afflict 1 in 5 people, it is essential to educate the next generation about the brain.

- **Artificial Intelligence (AI)** is today synonymous with **artificial neural networks**. These AIs can perform feats of classification, prediction and control we previously thought only humans could perform, such as generating images, controlling robots, conversing and playing video games. Familiarity with AI is becoming essential in many careers.

- **Neural modeling** is an important area of both neuroscience and AI research and is well represented within the Next Generation Science Standards (NGSS). **Neurorobots** make it easy to construct computational neuroscience models to control mobile robots in the classroom.

**SpikerBots** are educational neurorobots - robots controlled by neural networks running on laptops**.** SpikerBots allow teachers with no background in neuroscience or programming to run classroom labs with neural networks and robots, and help students investigate how brains process information, make decisions and navigate the world.

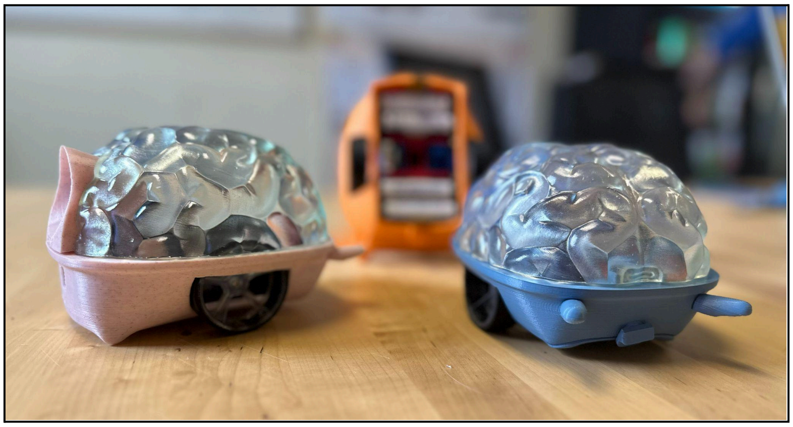

This **SpikerBot Primer for Teachers** is a guide to teaching neuroscience and artificial intelligence with robots. It is designed to provide teachers with neuroscience background knowledge, SpikerBot user instructions and teaching strategies. The primer should be used in combination with the SpikerBot **Lesson Plans**, which include lesson-specific goals, concepts, skills, exercises with solutions and student handouts.

This Primer is divided into three sections:

1. **Basics of Biological and Artificial Neural Networks**
2. **Neurorobot Basics: Using the Robot and the App**
3. **Teaching with SpikerBots**

### 1. Basics of Biological and Artificial Neural Networks

A **brain** is a **biological neural network** that allows an animal to see, hear, feel, move, find resources, plan and stay safe. The brain is made up of tiny cells called **neurons**, which are connected by **synapses**. In the human brain there are more than 85 billion neurons and each neuron is connected to thousands of other neurons.

Neurons have a cell body with dendrites and **axons**. Dendrites receive signals from sensors (e.g. eyes, ears, whiskers) and from other neurons. The cell body integrates these signals and generates electrical output signals called **action potentials** that travel via the axon to the synaptic contacts, which can **stimulate** or **inhibit** their targets. The brain controls behavior by sending action potentials to muscles. For example, when you want to walk, your brain sends signals to your leg muscles to make them move.

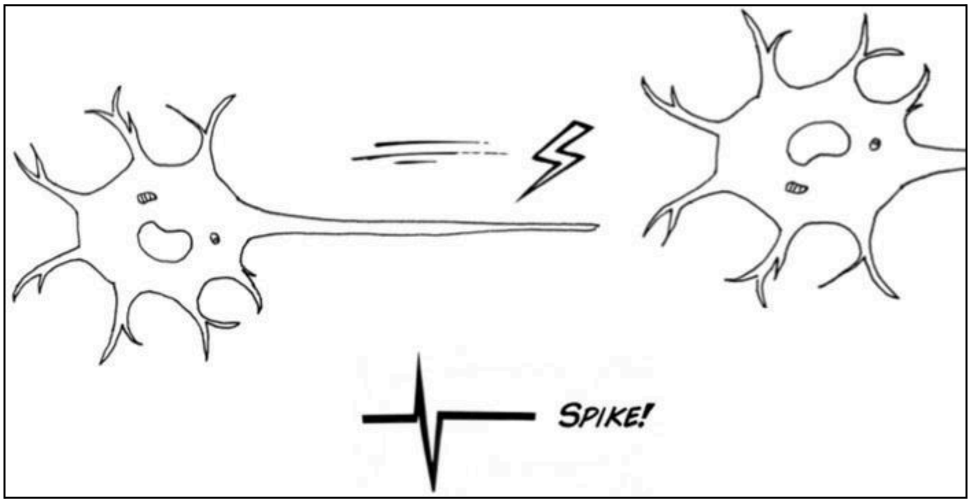


The **function** (behavior) of a neural network depends on its **structure** (connectome) - the types of neurons the network is made of and the way they're connected. Neurons can be quiet, sporadically active or bursting with activity. Synaptic connections can be short and simple, or winding and complex. In Lesson 1, students will design brains in which neurons monitoring one side of the visual field drive the wheel on the opposite side of the brain forward. This midline-crossing connectivity is common in nature and allows even very small brains, like those of newborn fish, to approach prey.

To decide which action to perform at any given moment, your brain relies on a structure called the **basal ganglia.** The basal ganglia receives information from the cortex and sends action selection decisions back to the cortex in a loop. The basal ganglia exerts a powerful inhibitory influence on many neural networks associated with specific behaviors. By selectively releasing individual networks from inhibition, while keeping many others suppressed, the basal ganglia effectively chooses which of many possible actions to perform in any given moment. In Lesson

2, students construct brains that use a computational model of the basal ganglia to select between different neural networks and their associated behaviors.

**Artificial neural networks** are computer algorithms that mimic some of the ways the human brain works. Using enormous volumes of training data and an algorithm called backpropagation, artificial neural networks can be trained to accomplish tasks that were once only possible for humans, like understanding a visual scene, playing a video game, drawing images, composing music and conversing. This is what we call **artificial intelligence** (AI) today. While there are clear differences between biological and artificial neural networks, both are made of neurons connected by synapses whose strengths (weights) can adapt and learn.

Neural networks have two types of learning. One is called **associative learning**, where the network finds patterns in large volumes of complex data. Neural networks are really good at this and can easily learn to identify recurring objects, landmarks, voices or faces. In Lesson 3, students will train neural networks to identify visual landmarks that help the robot navigate its environment. The other type of learning is called **reinforcement learning**, where a network learns, through trial and error, how to perform a task, like playing a new game. Neural networks are great at this type of learning too. In Lesson 3, students train neural nets to navigate between local landmarks to reach a target.

### 2. SpikerBot Basics

The **SpikerBot robot** is a neurorobot designed for the science classroom. It uses a camera to see, a microphone to hear, a speaker to communicate, and motors to move around. It maintains a permanent WiFi connection with the **SpikerBot app**, which runs its neural networks.

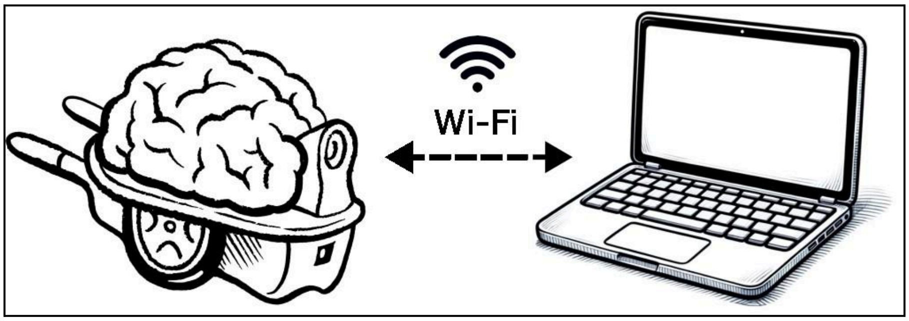


#### Installing the App on a Windows PC

- Download the repository at github.com/BackyardBrains/NeuroRobot
  - To download, click on the green 'Code' button, then 'Download zip'

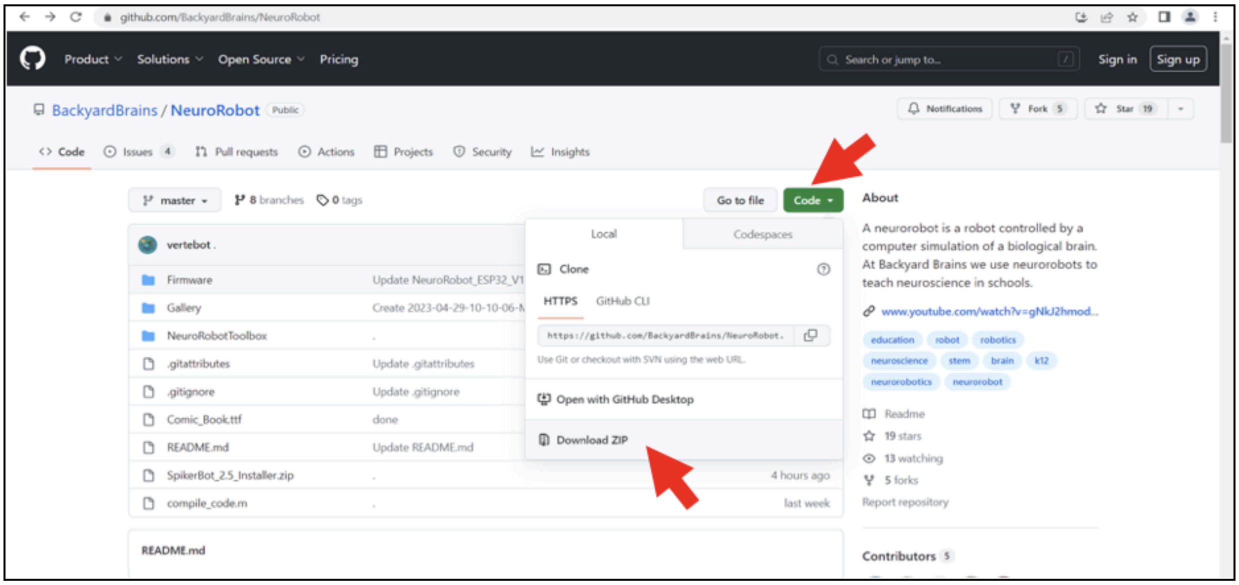


- Unpack the 'SpikerBot_Installer.zip' file inside the 'NeuroRobot-master' folders.
  - Click on the downloaded file at the bottom of your browser **OR** go into the 'Downloads' folder on your computer to locate the downloaded zip file.

- Run 'SpikerBot_Installer.exe' and follow the prompts on the window that pops up.

- ○ Because this is an unsigned app, you will need to navigate a blue warning message to begin the installation. If you see a ‘Windows protected your PC’ box pop up, click on ‘More info’, then the ‘Run anyway’ button.

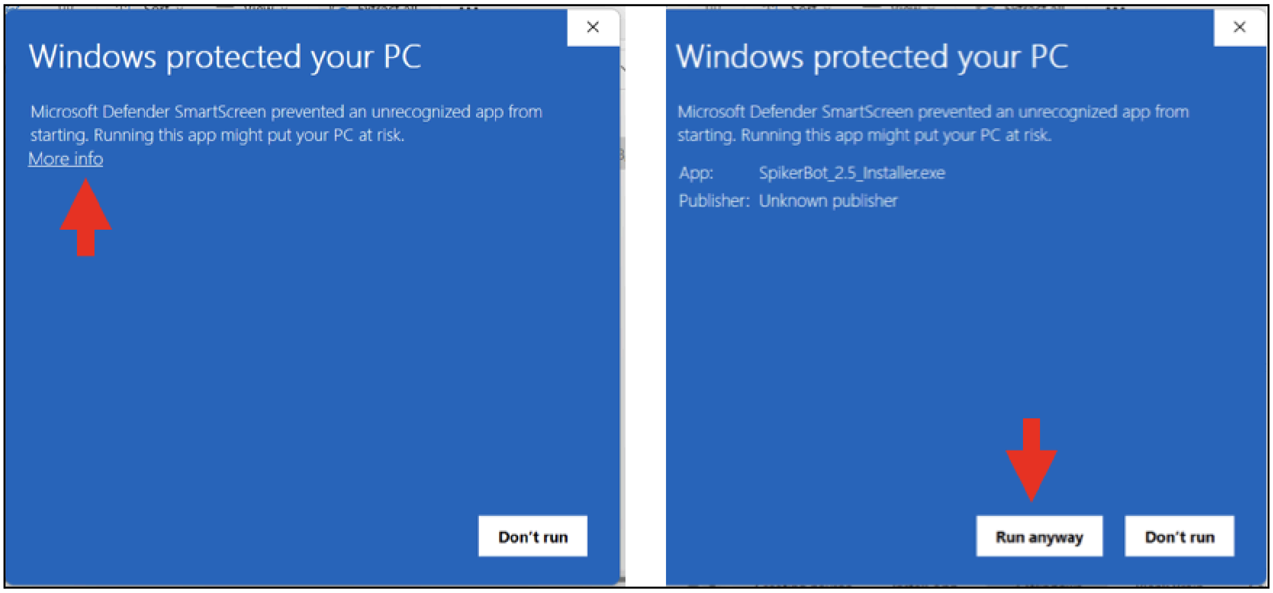


- Note the location of the destination folder of the SpikerBot App, so you can open it after installation.

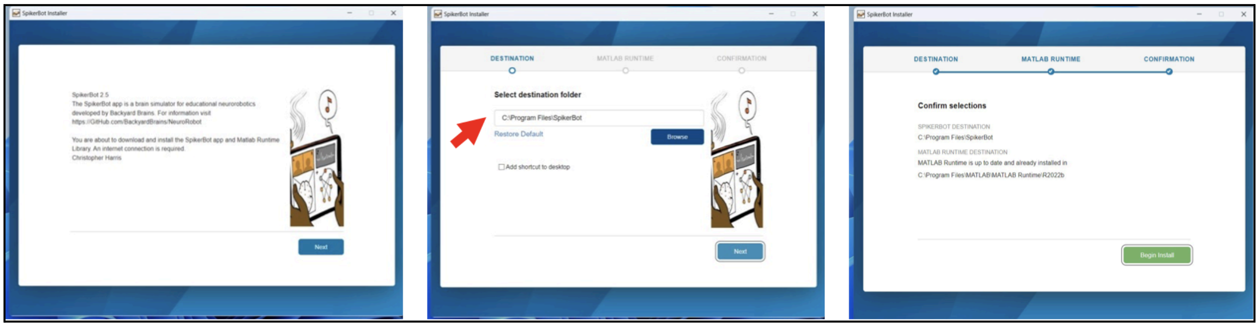


#### Quick Start - How to Start the SpikerBot Robot and App

1. The power switch is located on the underside of the neurorobot. Use it to turn the robot on.
2. Connect your computer to your robot’s WiFi network by accessing your computer’s WiFi settings and selecting the WiFi network name written on the robot (e.g. ‘NeuroRobot_400640’). Once connected, it will say "No internet access".
3. Start the SpikerBot app and click Connect. The button should turn green. This process may take up to 30 s depending on your laptop.

4. Select a brain from the list of brains by clicking on a name so it becomes highlighted. Alternatively, create and name a new empty brain. All other settings are default.
5. Once a brain is selected, click Runtime to allow the SpikerBot to operate. The Runtime button will turn green and then the Runtime mode window will open. This process may take up to 30 s.
6. Enter design mode by clicking Design to modify the brain.

#### How the App Works

##### <u>Modes of operation</u>

The SpikerBot app has 3 modes of operation: **Startup**, **Runtime** and **Design**.

**Startup** is the main menu. This is where you select which brain you want to work with, and make any special changes to settings when specified in lesson handouts (usually not needed).

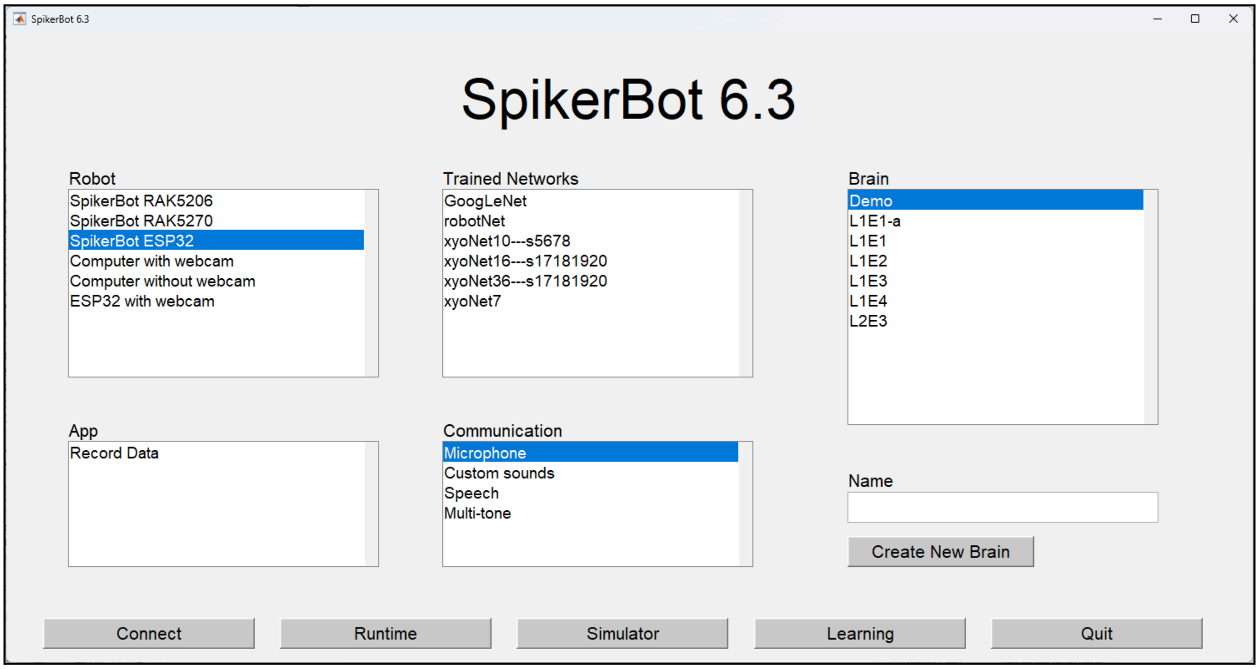


**Runtime** is the brain simulation engine where you can see what the brain is observing, hearing and doing.

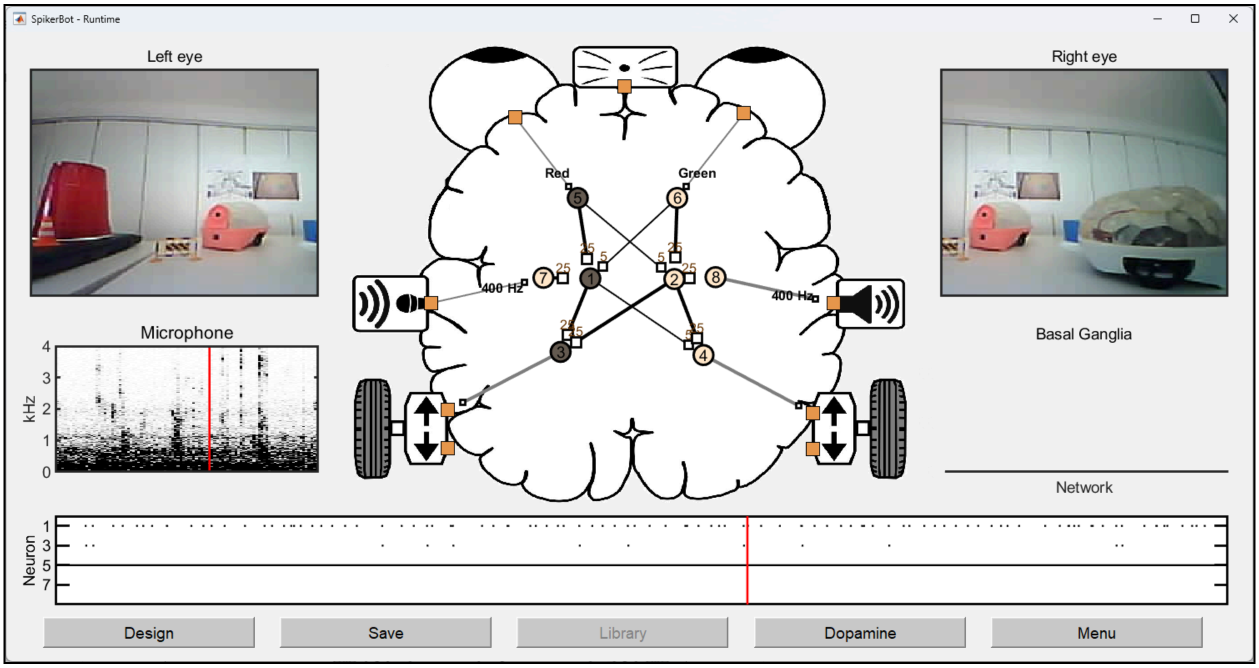


**Design** is where you modify the structure of the brain. You can click anywhere in the brain-shaped area to add a neuron or synapse.

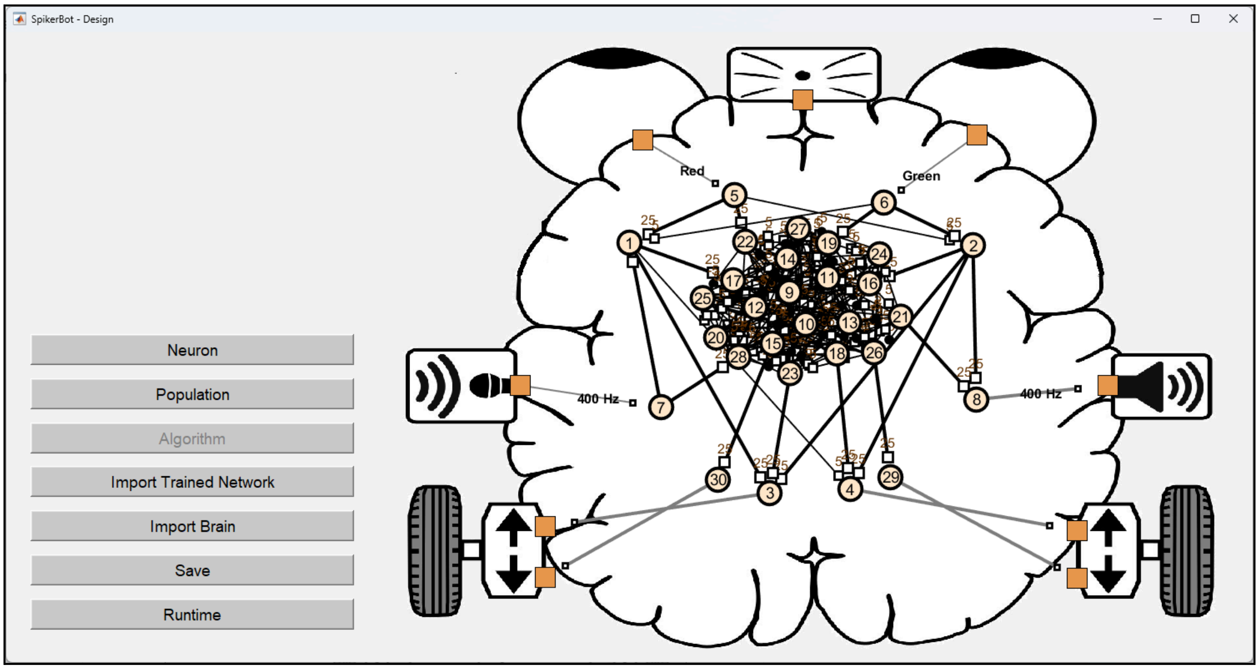


The app collects camera and microphone data continuously. Within this data, it can detect simple features such as color and pitch, and complex data such as objects and words. Below is a labeled diagram of the sensors (outlined green) and targets/effectors (outlined purple) you can connect to in design mode.

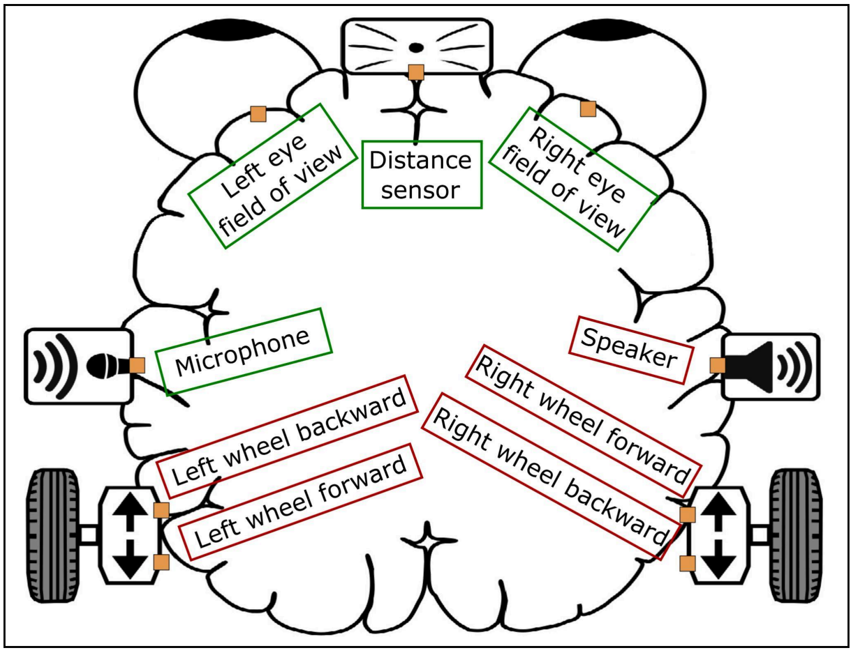


*Sensors & effectors in Design Mode*

**Switching between modes of operation**

- **Main Menu to Runtime mode:** Create a new brain or select an existing brain. Click the 'Runtime' button on the bottom left of the screen. The 'Runtime' button will turn green before entering Runtime mode.
- **Runtime mode to Design mode:** Click on the 'Design' button on the bottom left of the screen.
- **Design mode to Runtime mode:** Click 'Save' to save changes to brain design, then 'Runtime' on the bottom left of the screen.
- **Runtime mode to Main Menu:** Click 'Menu' on bottom right of screen.

**Deleting Brains from the app**

- Locate and enter the 'Documents' folder on your computer. Then enter the 'MATLAB' folder, then the 'Brains' folder.
- Delete the MATLAB file (.mat) for any brain(s) you would like to remove from the app.
- Upon reentering the SpikerBot app main menu, the deleted brain(s) will not appear in the Brains list.

**Object Detection via GoogLeNet**

- To enable the robot to detect some common household objects (such as cups, shoes, laptops), you can select 'GoogLeNet' under 'Trained Networks' in the Main Menu before running a brain.

#### Neurons

*The order in which components are connected matters!*

**Create, move, and delete a neuron**

- **Create neuron:** click anywhere in the brain and select type of neuron
- **Move neuron:** click on the neuron, then 'Move', click on new location in the brain
- **Delete neuron:** click on neuron, then 'Delete'

**Changing neuron type**

The simulated neurons are designed to model the spiking (firing) of biological neurons (for details, see Izhikevich, 2003). Thus, they can be quiet or fire regularly or in bursts, and can respond in different ways to synaptic inputs. Each neuron has 4 parameters (a-d) that can be adjusted to change how the neuron communicates:

- a - parameter regulates how quickly neurons recover from spikes
- b - parameter regulates spontaneous activity (firing rate)
- c - parameter modulates the duration of bursts
- d - parameter regulates the transition from bursting to spiking

In the SpikerBot app, there are 5 neuron types with pre-configured parameter settings (Quiet, Occasionally Active, Highly Active, Spontaneously Bursting, and Bursting when Stimulated). To change neuron type after a neuron is created, click on the neuron, then 'Properties' in the menu. Select the type of neuron you would like the neuron to be and click 'Confirm'.

#### Synapses

**Create and delete a synapse**

- **Create a synapse/extend an axon & modify synaptic weight:**
    - **Sensor to neuron:** click on orange square by sensor, click on neuron, select visual/distance/frequency preference depending on the input sensor (eyes, distance sensor or microphone), then 'Confirm'
    - **Neuron to neuron:** click on the first neuron, then 'Axon', then the second neuron. Choose excitatory or inhibitory. Change the synaptic weight if desired, then 'Confirm'
    - **Neuron to target/effector:** click on neuron, then 'Axon', then orange square by the target/effector (motor/wheel or speaker). Change the synaptic weight if desired, then create synapse (motor or sound output synapse depending on the effector).

- **Delete an axon:**
    - **Sensor to neuron OR neuron to target/effector:** repeat the same process as to create an axon, but in the final step click 'Delete synapse', then 'Confirm'

- **Neuron to neuron:** repeat the same process as to create an axon, then change synaptic weight to 0, then 'Confirm'

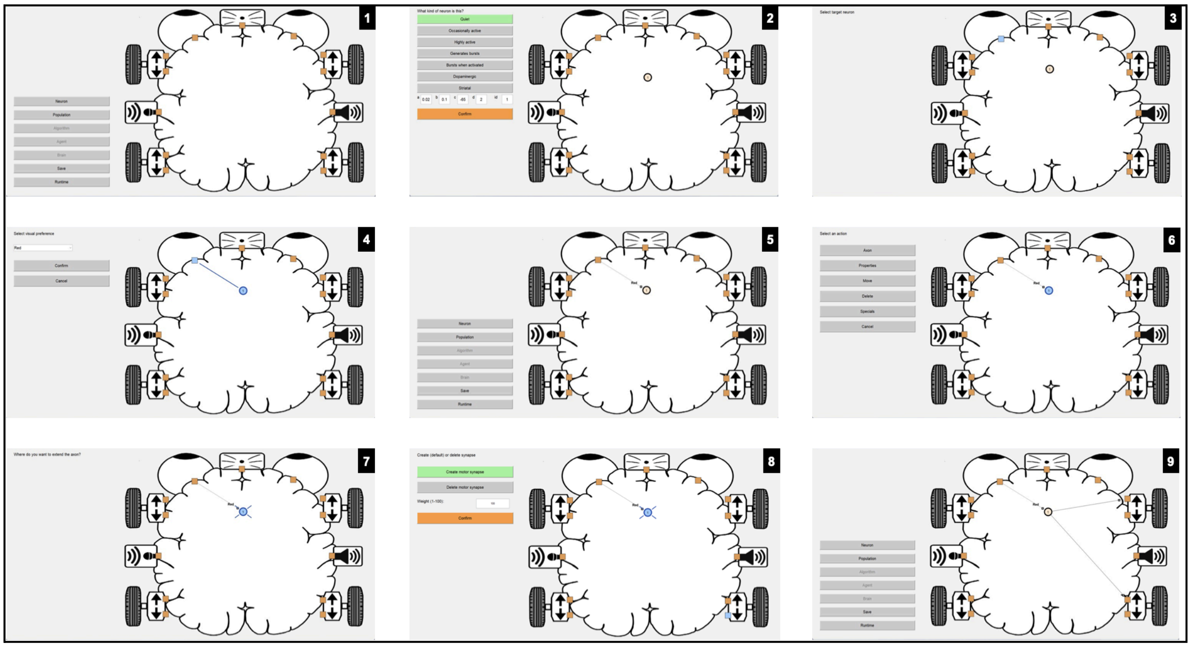

*The example above shows how to create a neuron, connect a sensor (camera - left eye) to the neuron, and then connect the neuron to a target/effector (motors - right side wheels forward).*

**Adjusting Synaptic Strength/Weight**

Synaptic connections between neurons have a strength ("weight") ranging from -100 to 100 mV. Every time a neuron fires a spike, the weight of the synapse is applied to the receiving ("postsynaptic") neuron's membrane potential. The membrane potential represents the final integration of all signals arriving at a neuron's cell body. To reliably trigger a spike in its target, a synapse should have a weight of 30 mV or more.

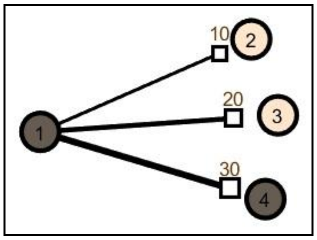


#### Basal Ganglia Model

The SpikerBot app includes a model of the basal ganglia's action selection process. Each neuron is assigned a basal ganglia network ID and each ID is associated with a dynamic **motivation** value that reflects the network's likelihood of being **selected**, i.e. win the brain's continuous "winner-take-all" competition. **Only neurons with a currently selected network ID can fire spikes.** ID 1 is exempt from selection, to allow for uninterrupted sensory input and activity in parts of the brain not connected to the basal ganglia.

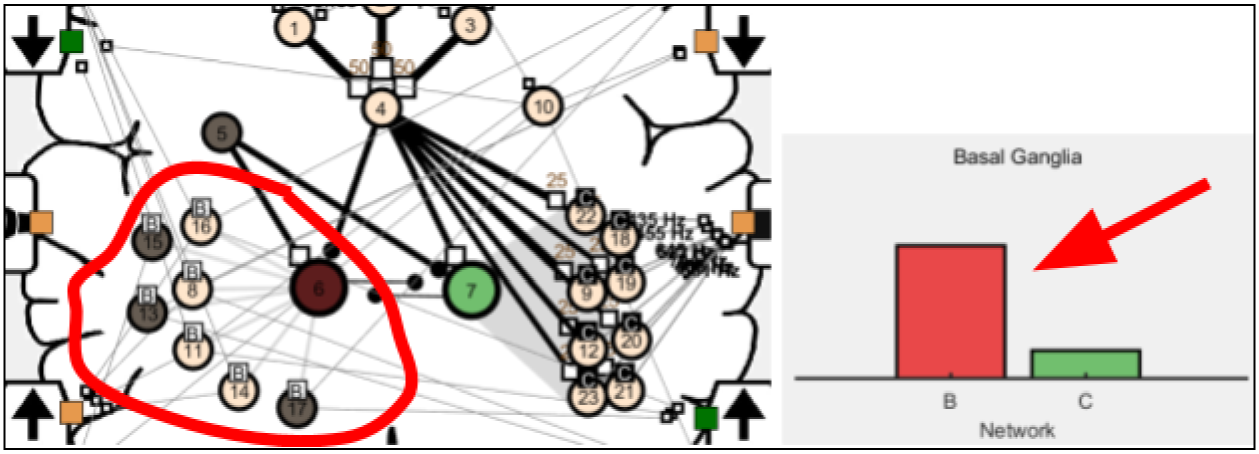


*Action selection in the basal ganglia. The left network (B) is currently selected (red circle, arrow). Some neurons in the network are generating action potentials (darker cell bodies). Neurons in the right network (C) remain inhibited by the basal ganglia and unable to generate spikes.*

The basal ganglia receives inputs from all over the brain, and different inputs promote or inhibit the selection of different networks. The basal ganglia's input neurons are called "**striatal**" neurons. They can be used to integrate sensory and other neuronal inputs intended to modulate the brain's motivation to select a specific network in particular situations. Synaptic inputs onto a striatal neuron increase the motivation of that neuron's network. In Lesson 2, students use striatal neurons to motivate a brain to explore in the absence of sensory input, and to stop exploring when the target is found.

### 3. Teaching with Neurorobots

#### Classroom Setup

Divide your students into groups of 2-4 students per group. Provide each group with a laptop, a robot and a lesson handout. Optionally, you can start and connect each robot-laptop pair before the class starts.

#### Troubleshooting

If the Connect or Runtime buttons fail to turn green, check the app's output window for errors. Check that the robot is on and that the laptop is connected to the correct robot's WiFi. WiFi connections sometimes fail unexpectedly and need to be manually restored.

If the robot is not detecting a specific color, check the camera feed in Runtime mode to see what the robot is 'seeing'. The shade of the color and lighting of the environment may be affecting the robot's ability to detect the color. If the lighting is bright, but the robot is still having difficulty recognizing a color, try pulling up an image of that color on your phone and test a few different shades.

#### Standards Alignment

Neuroscience is a new discipline that has yet to be fully integrated into state standards such as the NGSS. However, the SpikerBot curriculum has a contemporary emphasis on active learning and is well-aligned with NGSS Core Ideas LS1.A: Structure and Function and ETS1.B: Developing Possible Solutions to Real-world problems. Specifically, the curriculum contains brains that demonstrate important brain-behavior (LS: structure-function) relationships, and other brains that are impaired in some way and in need of an environmental or "surgical" solution (ETS: solutions).
