## Supplementary material for "Building Brains for Robots: A Hands-On Approach to Learning Neuroscience in the Classroom": Survey

### NEUROROBOT WORKSHOP SURVEY

Thank you for taking part in this neurorobot workshop. This survey is designed to help us at Backyard Brains measure how much you learn by taking part in the workshop. You will be given a similar quiz at the end of the workshop and we'll compare the results. But don't stress! Your teachers will not see your results and they will not affect your grades. If there's a question you can't answer, just leave it blank. We will not show your survey responses to anyone else - all results will be anonymized.

**Name** ______________________________________________

**Gender** ____________________ **Ethnicity** ____________________

#### NEUROSCIENCE QUIZ

**Describe three types of neuron with different spiking patterns**

______________________________________________

______________________________________________

______________________________________________

**How does the basal ganglia control behavior?**

______________________________________________

______________________________________________

______________________________________________

**Describe how neural networks learn new things**

______________________________________________

______________________________________________

______________________________________________

#### SCIENCE ATTITUDES

The following statements relate to descriptions of yourself in the science fields. Please indicate your level of agreement or disagreement with each statement.

| | Strongly Agree | | | Neutral | Disagree | | |
|---|---|---|---|---|---|---|---|
| | 1 | 2 | 3 | 4 | 5 | 6 | 7 |
| I am confident I can recognize the science question that underlies a newspaper report on a neuroscience issue | | | | | | | |
| I am confident I can explain basic neuroscience concepts we observe in the real world | | | | | | | |
| I am confident I can describe the role of key elements of neuroscience processes | | | | | | | |
| I am confident I can understand complex material in neuroscience | | | | | | | |
| I am confident I can complete the activities in the neurorobot workshop | | | | | | | |
| I am confident I can interpret basic information presented in neuroscience | | | | | | | |
| I am confident I can master the concepts in the neurorobot workshop | | | | | | | |

The following statements relate to your self-confidence in science. Please indicate your level of agreement or disagreement with each statement.

| | Strongly Agree | | | Neutral | Disagree | | |
|---|---|---|---|---|---|---|---|
| | 1 | 2 | 3 | 4 | 5 | 6 | 7 |
| Learning advanced science topics would be easy for me | | | | | | | |
| I can usually give good answers to test questions on science topics | | | | | | | |
| I learn science topics quickly | | | | | | | |
| Science topics are easy for me | | | | | | | |
| When I am being taught science I can understand the concepts very well | | | | | | | |
| I can easily understand new ideas in science | | | | | | | |
| Compared to others my age I am good at science classes | | | | | | | |
